# Oxidized TRX1 and DPP8/9 stabilize the MIA40-AIFM1 complex during intrinsic apoptosis

**DOI:** 10.64898/2026.10.26.761918

**Authors:** Lydia P. Tsamouri, Michael B. Geeson, Marianna Teplova, Buren Li, Zhencheng Lai, Andrea V. Tellez, Daniel P. Ball, Arpita Chakravarti, Sagnik Sen, M. Jason de la Cruz, Daniel A. Bachovchin

## Abstract

Mammalian cells execute diverse death programs with distinct immunological consequences. Mitochondrial outer membrane permeabilization (MOMP) drives typically immunologically silent intrinsic apoptosis by releasing cytochrome c and other proapoptotic proteins into the cytosol. Here, we show that MOMP also engages two established repressors of the pyroptosis-inducing NLRP1 inflammasome, oxidized thioredoxin-1 (oxTRX1) and catalytically active DPP8/9, in a coordinated redox and proteostasis program. Immediately after MOMP, oxTRX1 forms a disulfide with MIA40 Cys4 at the MIA40–AIFM1 interface. Subsequently, DPP8/9 cleave AK2, targeting this competing AIFM1 ligand for degradation. Biochemical reconstitution and cryo-EM reveal that oxTRX1 promotes AIFM1 dimerization and preferentially stabilizes MIA40–AIFM1 over AK2–AIFM1. Thus, redox-dependent complex assembly and selective proteolysis converge to favor MIA40-bound AIFM1 after MOMP. These findings suggest how post-MOMP remodeling could reinforce NLRP1 restraint during apoptosis and how disruption of this program could trigger pyroptosis.

## Introduction

Mammalian cells can die in a remarkable variety of ways^1,2^. Intrinsic apoptosis is typically nonlytic and immunologically silent. During intrinsic apoptosis, developmental cues or cellular stress activate BAX and BAK, causing mitochondrial outer membrane permeabilization (MOMP) and cytochrome c release. Cytosolic cytochrome c then promotes apoptosome assembly and activation of the initiator caspase-9, which in turn activates executioner caspases-3 and -7 to drive orderly cellular dismantling^3,4^. By contrast, pyroptosis is highly inflammatory and lytic^5^. Pyroptosis is initiated when specific stress or danger signals trigger pattern-recognition receptors (PRRs) to assemble into multiprotein signaling complexes called inflammasomes, leading to inflammatory caspase (caspase-1/4/5) activation, gasdermin D-mediated membrane rupture, and inflammatory cytokine release. Despite substantial progress in defining programmed cell death pathways over the past three decades, the signals that engage particular cell death programs, as well as the mechanisms through which distinct death pathways communicate with one another, remain incompletely understood ^1,2,6,7^.

NLRP1 is a cytosolic PRR that can form an inflammasome (**Fig. S1A**)^8,9^. NLRP1 is normally restrained through direct interactions with the oxidized form of thioredoxin-1 (oxTRX1)^10–12^ and with the catalytically active forms of the closely related serine proteases DPP8 and DPP9 (DPP8/9) ^13–15^. Both regulatory mechanisms are broadly conserved: oxTRX1^10,11^ and DPP8/9^16,17^ bind and repress functional human, rat, and mouse NLRP1 alleles, and DPP9 also binds and represses zebrafish NLRP1^17^. Accordingly, synthetic DPP8/9 inhibitors such as Val-boroPro (VbP) activate NLRP1 inflammasomes in all species tested^16,18–20^. However, the relevant substrates and regulators of DPP8/9, the physiological relationship between DPP8/9 and oxTRX1, and the endogenous danger signal that overcomes these NLRP1 restraints all remain unknown.

Notably, TRX1 directly connects NLRP1 to the cytosolic redox state. In the canonical thioredoxin cycle, Cys32 of reduced TRX1 (redTRX1) attacks a substrate disulfide, forming a transient intermolecular disulfide. Cys35 then resolves this intermediate, producing a reduced dithiol in the substrate and a Cys32-Cys35 intramolecular disulfide in ox-TRX1. Thioredoxin reductase-1 (TXNRD1) subsequently uses NADPH to regenerate redTRX1. Distinct from this cycle, Cys32 of oxTRX1 forms a stable intermolecular disulfide bond with Cys427 of NLRP1 that represses inflammasome activation (**Fig. S1S**) ^10–12^. A sufficiently reducing environment could therefore disfavor oxTRX1–NLRP1 binding and activate NLRP1^10^. Our discovery that the redox sensor KEAP1 inhibits DPP9 suggests that a common redox event may regulate both NLRP1 restraints^21^.

Despite this progress, the physiological DPP8/9 substrates relevant to NLRP1 regulation have remained elusive ^9,19,22^. Notably, adenylate kinase 2 (AK2), a mitochondrial intermembrane space (IMS) enzyme that catalyzes the reversible transfer of phosphate groups among adenine nucleotides (AMP + ATP ↔ 2ADP)^23^, was identified as a potential DPP8/9 substrate by N-terminal proteomics in cells overexpressing DPP8/9^24^. However, because mature AK2 and DPP8/9 reside in the IMS and cytosol, respectively, it was unclear whether DPP8/9 encounter AK2 in intact cells. Riemer and colleagues subsequently proposed that mitochondrial import of AK2 competes with its processing by cytosolic DPP8/9, with cleavage exposing a destabilizing N-terminal residue that targets AK2 for N-degron-mediated proteasomal degradation^25,26^. This model suggests that DPP8/9 eliminate AK2 molecules that fail to enter the IMS efficiently, thereby limiting their cytosolic abundance. However, only small amounts of cytosolic AK2 accumulated following DPP8/9 inhibition, suggesting that this degradation pathway normally operates at low flux and leaving it unclear whether AK2 ever becomes a major physiological DPP8/9 substrate.

Because DPP8/9 processing competes with mitochondrial import, the MIA40 oxidative-folding pathway that imports AK2 is itself a determinant of AK2 stability (**Fig. S1B,C**). MIA40 and AIFM1 are the core components of this pathway^25,27–29^. MIA40 has an N-terminal β-hairpin-forming regulatory region, a redox-active CPC motif (residues 53– 55), and a disulfide-stabilized substrate-binding domain (**Fig. 1A**). AIFM1 contains an N-terminal mitochondrial-localization sequence and transmembrane domain (MLS-TMD) that anchors it to the inner mitochondrial membrane (IMM), and a C-terminal NADH- and FAD-dependent oxidoreductase domain that extends into the IMS (**Fig. 1A,B**). In the prevailing model, NADH binding promotes AIFM1 dimerization and displacement of its C-loop, exposing a docking surface for the MIA40 N-terminus (residues 1–30) (**Fig. 1B**)^30–32^. AIFM1 binding relieves MIA40 autoinhibition by displacing the N-terminal β-hairpin from the substrate-binding domain and thereby enhances oxidative folding activity^32^. During import, reduced and unfolded MIA40 substrates, including MIA40 itself and AK2, engage the MIA40 substrate-binding domain, enabling the CPC motif to transfer a disulfide that promotes oxidative folding and traps the protein within the IMS (**Fig. S1B,C**). The MIA40 CPC motif is subsequently reoxidized by the sulfhydryl oxidase ALR^33^ (**Fig. S1S**). Intriguingly, recent studies found that AK2 and MIA40 bind the same surface on AIFM1, although the functional significance of this competition remains unclear ^34,35^.

**Figure 1.**
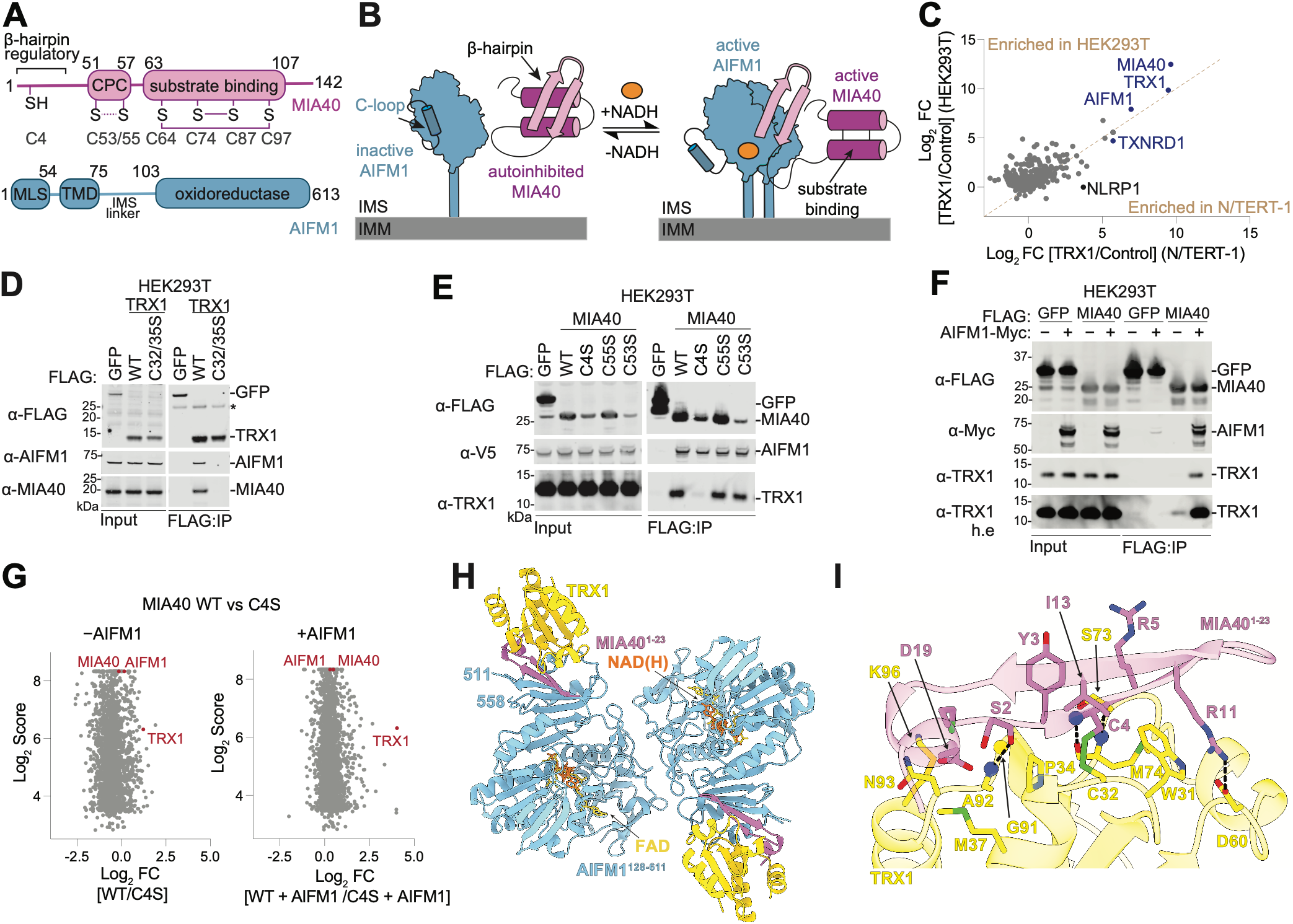
TRX1 forms a ternary complex with AIFM1 and MIA40. (**A**) Domain organization of MIA40 and AIFM1. MIA40 redox-active and structural disulfides are shown as solid and dashed lines, respectively. (**B**) Schematic of NADH-dependent AIFM1 dimerization and MIA40 activation. (**C**) FLAG–TRX1 IP-MS in HEK 293T and N/TERT-1 cells. (**D–F**) The indicated proteins were expressed in HEK 293T cells, IPed using anti-FLAG beads, and analyzed by immunoblotting. (**G**) FLAG–MIA40 IP-MS comparing WT and Cys4Ser MIA40 in the absence (left) or presence (right) of co-expressed AIFM1. (**H**) Cryo-EM structure of the AIFM1–MIA40^1–45^ fusion bound to TRX1 (PDB 38EA). (**I**) Close-up view of the TRX1–MIA40 interface. Immunoblots are representative of three or more independent experiments.

Collectively, DPP8/9 and oxTRX1 are now established as the key NLRP1 regulators^10–14,19,20^, but no physiological connection between them has been identified. Here, we show that MOMP, widely considered the commitment step of intrinsic apoptosis, initiates a coordinated redox and proteostasis program that brings these two NLRP1 restraints together. Following MOMP, oxTRX1 forms a disulfide with MIA40 Cys4 at the MIA40–AIFM1 interface and promotes AIFM1 dimerization, favoring the MIA40-bound state over the competing AK2–AIFM1 complex. DPP8/9 subsequently degrade AK2, reinforcing MIA40 occupancy. Thus, oxTRX1 and DPP8/9 are not only NLRP1 repressors, but also components of a common post-MOMP remodeling program. These findings suggest how post-MOMP remodeling could reinforce NLRP1 restraint during apoptosis and how disruption of this program could favor inflammatory cell death when apoptotic execution fails.

## Results

### TRX1 forms a ternary complex with AIFM1 and MIA40

We hypothesized that oxTRX1 regulates proteins beyond NLRP1, and that identifying these targets could reveal the danger state that activates NLRP1. We therefore expressed FLAG-tagged TRX1 in HEK 293T cells and N/TERT-1 keratinocytes and performed anti-FLAG immunoprecipitation followed by quantitative mass spectrometry (IP-MS) (**Fig. 1C**). As expected, TRX1 and TXNRD1 were recovered in both cell lines, whereas NLRP1 was detected only in N/TERT-1 keratinocytes, consistent with its absence in HEK 293T cells ^10,11^. Notably, AIFM1 and MIA40 were among the most strongly enriched proteins in both cell lines. Although TRX1 associations with AIFM1 and/or MIA40 have been reported in two previous IP-MS studies ^36,37^ and in the BioPlex Interactome^38^, these interactions are difficult to reconcile with their subcellular localization: TRX1 is cytosolic, whereas AIFM1 and MIA40 reside in the IMS. Their molecular basis and functional significance have remained largely unexplored.

We first asked whether any specific cysteines were required for these interactions. We found that TRX1 Cys32Ser and TRX1 Cys32Ser/Cys35Ser failed to detectably bind either MIA40 or AIFM1, and TRX1 Cys35Ser showed markedly reduced binding to both proteins (**Fig. 1D; Fig. S1S**). Thus, the TRX1 active site mediates binding to MIA40 and AIFM1. We next examined the reciprocal MIA40 determinants. Intriguingly, MIA40 Cys4, which lies within the β-hairpin that binds to AIFM1 (**Fig. 1A**), is conserved across metazoans but has no known function (**Fig. S1S**). We found that MIA40 Cys4Ser failed to bind TRX1 but retained binding to AIFM1 (**Fig. 1E**), demonstrating that Cys4 is selectively required for TRX1 recruitment rather than for formation of the MIA40–AIFM1 complex^39^. By contrast, mutation of the conserved redox-active cysteines in the MIA40 CPC motif, Cys53 and Cys55, to serine did not disrupt binding to either TRX1 or AIFM1 (**Fig. 1E**). Consistent with a redox-sensitive TRX1 interaction, DTT reduced TRX1 association with WT and Cys53Ser/Cys55Ser MIA40 without substantially affecting AIFM1 binding (**Fig. S1S**). More broadly, WT and cysteine-mutant MIA40 proteins, including cysteine-free (CysSer7) MIA40, all co-immunoprecipitated AIFM1, confirming that MIA40 cysteines are dispensable for AIFM1 binding^39^.

Notably, recovery of endogenous TRX1 in MIA40-FLAG IPs was also enhanced by AIFM1 co-expression (**Fig. 1F**), consistent with the AIFM1-bound MIA40 β-hairpin forming a composite surface for TRX1 recruitment. Similarly, quantitative MS comparison of wild-type (WT) and Cys4Ser MIA40-FLAG IPs showed that AIFM1 co-expression markedly increased recovery of TRX1 with WT, but not Cys4Ser, MIA40 (**Fig. 1G; Fig. S1S**). These findings identify MIA40 Cys4 as a selective docking site for TRX1 within the AIFM1-bound complex. Finally, Blue Native PAGE (BN-PAGE) revealed co-migration of AIFM1, MIA40, and TRX1 in a higher-molecular-weight complex, both in IPs from cells coexpressing AIFM1 and MIA40 and after co-incubation of purified recombinant AIFM1, MIA40, and TRX1 (**Fig. S1H, I**). Together, these biochemical and proteomic data indicate that TRX1, MIA40, and AIFM1 form a ternary complex dependent on TRX1 Cys32 and MIA40 Cys4.

We next determined the structure of this complex. We purified a TRX1 variant in which the three noncatalytic cysteines, Cys62, Cys69, and Cys73, were mutated to serine to prevent self-oligomerization *in vitro*^*40*^. This protein was then incubated with NADH and the previously described AIFM1-MIA40 fusion protein comprising AIFM1 residues 104–613 linked to MIA40 residues 1–45^32^. TRX1 co-eluted with the AIFM1–MIA40 fusion during size-exclusion chromatography (SEC, **Fig. S1S**). The 2.8 Å consensus cryo-EM reconstruction revealed a symmetric dimeric assembly containing two copies each of AIFM1, MIA40, and TRX1 (**Fig. 1H; Fig. S1S**). Each AIFM1 protomer adopted the dimeric, C-loop-displaced conformation with clear density for a bound dinucleotide cofactor and for the MIA40 β-hairpin^32,34^ (**Fig. 1H**). TRX1 bound the MIA40 β-hairpin on the surface of AIFM1, with density around the TRX1 active site consistent with a mixture of intermolecular MIA40 Cys4–TRX1 Cys32 and intramolecular TRX1 Cys32–TRX1 Cys 35 disulfide states (**Fig. S1S**). Backbone hydrogen bond interactions flanking the disulfide bond form an intermolecular β-sheet, supplemented by a salt bridge between TRX1 Asp60 and MIA40 Arg11 (**Fig. 1I**). Strikingly, this interface closely resembles the previously described TRX1–NLRP1 (PDB 7WGE)^11,12^ and TRX1–TXNIP (PDB 4LL1) ^41^ complexes (**Fig. S1S**). Thus, oxTRX1 engages the AIFM1-bound MIA40 N terminus through a Cys32–Cys4 intermolecular disulfide.

### MIA40 Cys4 forms an intermolecular disulfide bond with TRX1 Cys32

We next sought to directly detect a covalent adduct between TRX1 Cys32 and MIA40 Cys4. We adapted a thiol-trapping strategy previously used to detect the TRX1 Cys32–NLRP1 Cys427 conjugate^12^, in which samples are treated with Nethylmaleimide (NEM) to alkylate free thiols and prevent disulfide exchange, separated by nonreducing SDS-PAGE, and analyzed by immunoblotting (**Fig. 2A**). We first applied this strategy to purified AIFM1, MIA40, and TRX1 proteins and to the eluates from the MIA40-FLAG IP experiment described above (**Fig. 1F**). In both settings, we detected an AIFM1-dependent higher-molecular-weight species recognized by both MIA40 and TRX1 antibodies under nonreducing, but not reducing, conditions (**Fig. S2A,B**, cyan arrowheads). We next co-expressed WT or Cys4Ser MIA40 together with TRX1 and AIFM1 in HEK 293T cells and subjected the lysates to the same trapping protocol. As expected, WT MIA40, but not Cys4Ser MIA40, formed a DTT-sensitive conjugate with TRX1 in the presence of co-expressed AIFM1 (**Fig. 2B**, cyan arrowheads). These results demonstrate that MIA40 Cys4 and TRX1 form an AIFM1-dependent disulfidelinked conjugate.

**Figure 2.**
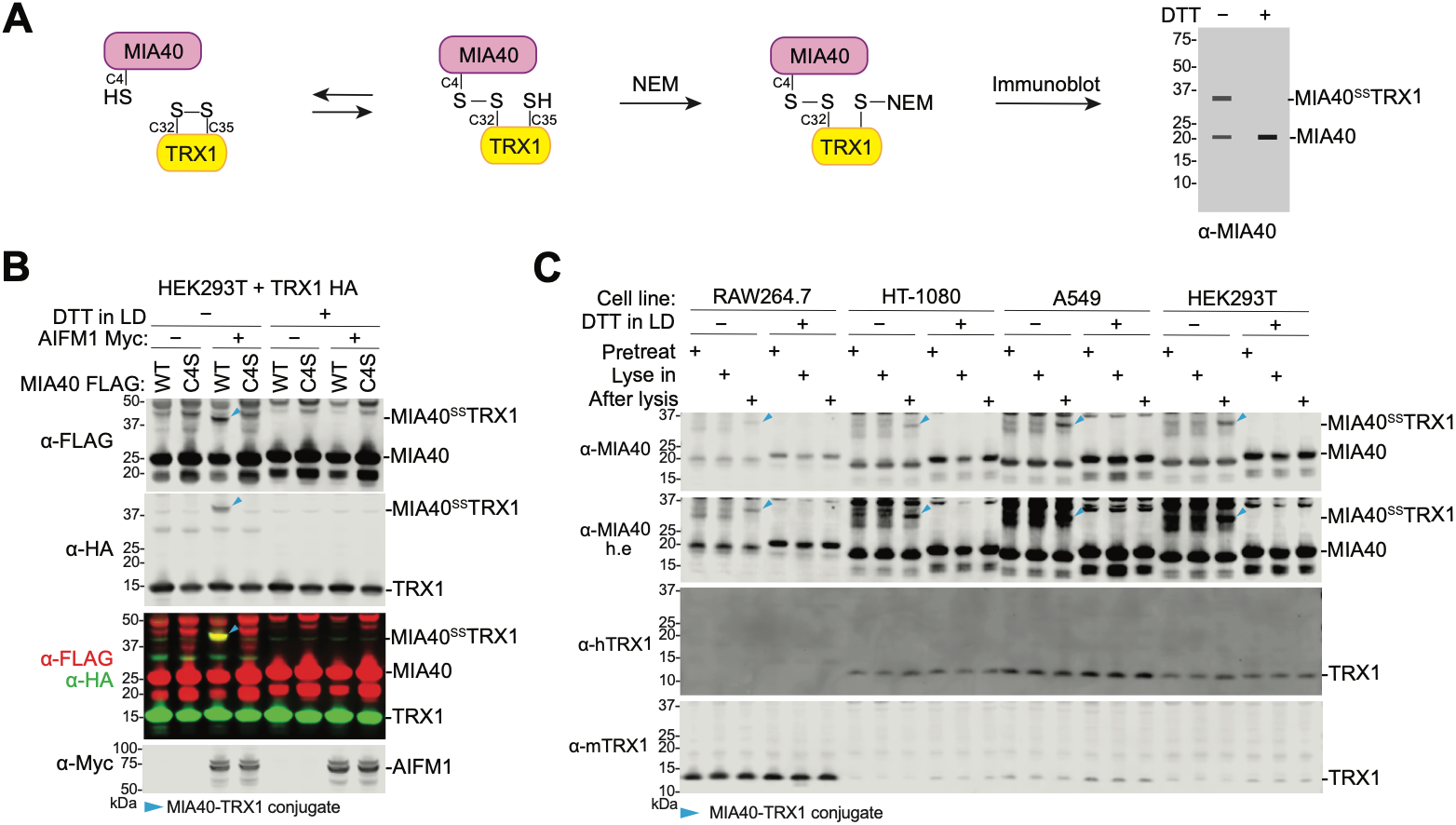
MIA40 Cys4 forms an intermolecular disulfide bond with TRX1 Cys32. (**A**) Schematic of the NEM-based thiol-trapping assay used to detect the MIA40–TRX1 conjugate by nonreducing immunoblotting. (**B–C**) HEK293T cells expressing the indicated proteins (**B**) and the indicated cell lines (**C**) were subjected to the specified NEM (40 mM) thiol-trapping conditions (either before lysis, during lysis, or after lysis) and analyzed by immunoblotting under reducing or nonreducing conditions (LD, loading dye). Arrowheads indicate the MIA40–TRX1 conjugate. Immunoblots are representative of three or more independent experiments.

We next asked whether this conjugate could form between the endogenous proteins and whether it existed in intact cells before lysis. Mouse RAW 264.7 cells and human HT-1080, A549, and HEK 293T cells were treated with NEM before, during, or after lysis, and conjugate formation was assessed by reducing and nonreducing SDS-PAGE (**Fig. 2C**). In each cell line, the disulfide-linked MIA40 species was detected only when NEM was added after lysis, but not when NEM was added before or during lysis (**Fig. 2C**, cyan arrowheads). By contrast, the TRX1–NLRP1 disulfide-linked conjugate was readily detected when iodoacetamide or NEM was added before, during, or after lysis (**Fig. S2S**, cyan arrowheads). These findings suggest that cellular compartmentalization prevents MIA40, but not NLRP1, from accessing ox-TRX1 under basal conditions.

Endogenous TRX1 was not detected within the conjugate by immunoblotting (**Fig. 2C**), likely because of antibody limitations. To confirm the identity of the MIA40-conjugated protein, we expressed HA-tagged TRX1 and observed the predicted increase in the apparent molecular weight of the conjugate detected by MIA40 immunoblotting (**Fig. S2S**). Moreover, endogenous conjugate formation was severely diminished in *MIA40*- and *AIFM1*-deficient cells (**Fig. S2E,F**). Collectively, these results establish that MIA40 Cys4 forms an AIFM1-dependent, disulfide-linked conjugate with TRX1 Cys32 and that, under basal conditions, this conjugate is generated only after disruption of cellular compartmentalization.

### The MIA40–TRX1 conjugate forms after MOMP

We reasoned that MOMP, widely considered the commitment step of intrinsic apoptosis^4^, might permit conjugate formation in intact cells by disrupting the compartmental separation of TRX1 and MIA40 (**Fig. 3A**). To test this possibility, we treated RAW 264.7 and HT-1080 with a panel of intrinsic apoptosis-inducing agents and assessed MIA40– TRX1 conjugate formation. Staurosporine, raptinal, and combined treatment with either ABT-737 or ABT-263 plus S63845 induced PARP and CASP3 cleavage, markers of apoptosis, and, consistent with our hypothesis, also induced formation of the MIA40 conjugate (**Fig. 3B, Fig. S3S**). We used ABT-263 plus S63845, hereafter termed ABT/S6, in many subsequent experiments because this combination is a mechanistically defined method for rapidly inducing intrinsic apoptosis by inhibiting BCL2 family proteins^42^. Consistent with a requirement for MOMP, ABT/S6 and staurosporine induced substantially less CASP3 cleavage and MIA40 conjugate formation in BAX/BAK-deficient RAW 264.7 cells (**Fig. 3C, Fig. S3S**). By contrast, raptinal, which disrupts the outer mitochondrial membrane (OMM) independent of BAX/BAK^43^, induced both CASP3 cleavage and conjugate formation in BAX/BAK-deficient cells (**Fig. 3C**). ABT/S6 also induced MIA40 conjugate formation in immortalized N/TERT-1 keratinocytes (**Fig. S3S**), resting and activated primary human T cells (**Fig. S3S**), and primary human peripheral blood mononuclear cells (PBMCs) (**Fig. S3S**), demonstrating that this response occurs across diverse cell types and metabolic states. Pyroptotic or ferroptotic stimuli that do not induce MOMP failed to generate the conjugate (**Fig. S3F, G**).

**Figure 3.**
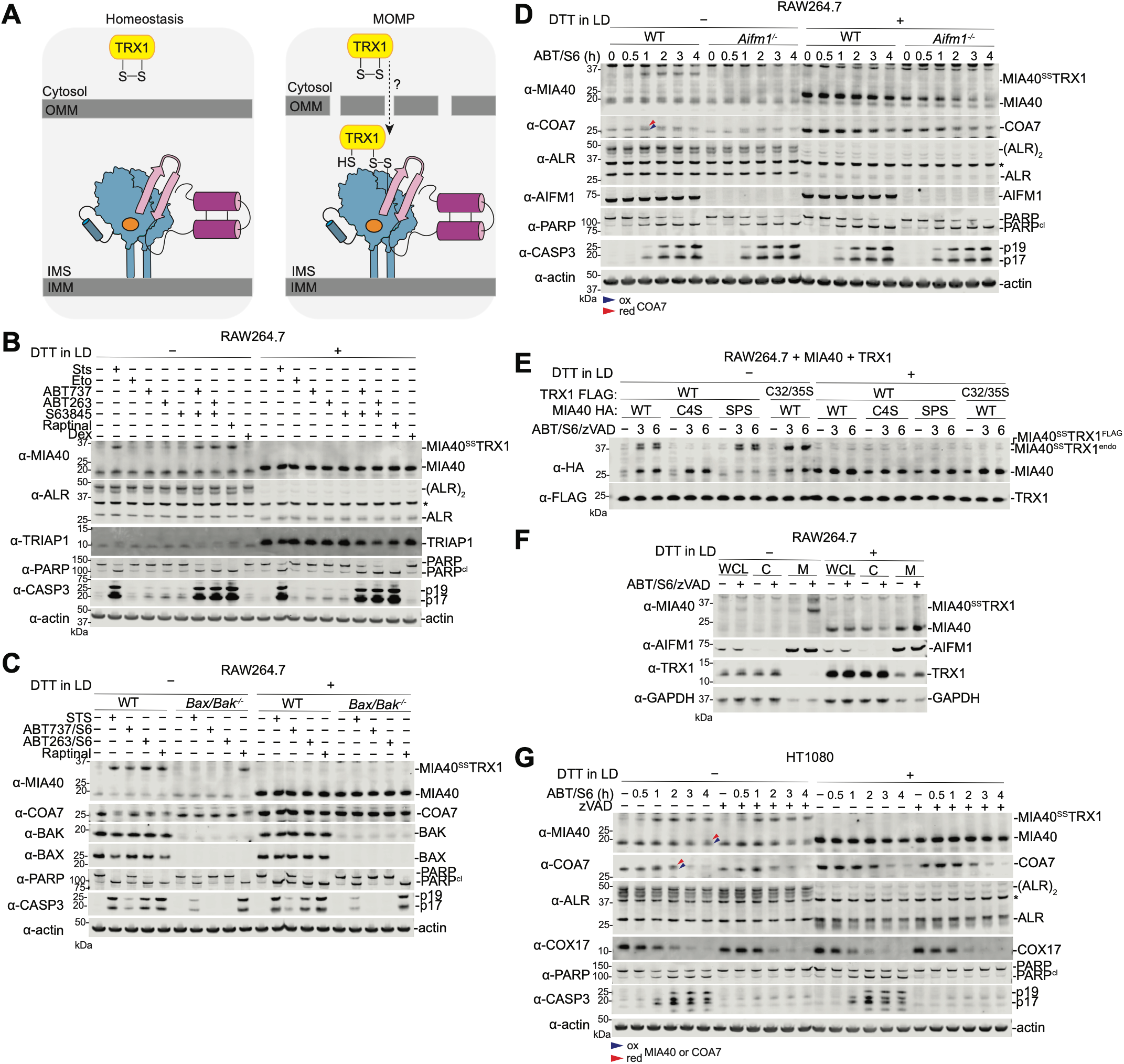
The MIA40–TRX1 conjugate forms after MOMP. (**A**) Proposed model of TRX1–MIA40–AIFM1 complex formation following MOMP. (**B– D**) RAW264.7 WT (**B**), BAX/BAK- (**C**), or AIFM1-deficient (**D**) cells were treated with the indicated compounds for 2 h (**B, C**) or for the indicated timepoints (**D**). (**E-G**) RAW 264.7 cells stably overexpressing the indicated MIA40 and TRX1 constructs (**E**) and HT-1080 cells (**G**) were treated for the indicated times in the presence or absence of zVAD, while RAW 264.7 cells were subjected to mitochondrial fractionation (WCL, whole cell lysate; cytosol, C; mitochondria, M) following ABT/S6 treatment for 2 h (**F**). Arrowheads in **D** and **G** indicate the reduced and oxidized forms of MIA40 and COA7. Staurosporine (10 μM), etoposide (20 μM), ABT-737 (10 μM), ABT-263 (10 μM), S63845 (1 μM), raptinal (10 μM), dexamethasone (50 μM), and zVAD (50 μM) were used as indicated. All cells were lysed in buffer containing 40 mM NEM and analyzed by immunoblotting under reducing or nonreducing conditions. Immunoblots are representative of three or more independent experiments. Asterisks denote non-specific bands.

We next verified that apoptosis-induced conjugate corresponded to the AIFM1-dependent MIA40–TRX1 complex. As expected, depletion of *AIFM1* (**Fig. 3D**) or *MIA40* (**Fig. S3S**) markedly reduced conjugate formation. Moreover, the ABT/S6-induced conjugate formed with ectopically expressed WT or Cys53Ser/Cys55Ser MIA40 and with WT TRX1, but not with Cys4Ser MIA40 or Cys32Ser/Cys35Ser TRX1 (**Fig. 3E**). Finally, subcellular fractionation and differential centrifugation showed that the ABT/S6-induced conjugate was enriched in mitochondrial (**Fig. 3F**) and insoluble fractions (**Fig. S3S**), respectively, consistent with formation at the inner mitochondrial membrane (IMM), where AIFM1 is anchored. Together, these results establish that intrinsic apoptosis induces formation of an AIFM1-dependent, disulfide-linked conjugate between MIA40 Cys4 and TRX1 Cys32 downstream of MOMP (**Fig. 3A**).

We next sought to position MIA40–TRX1 conjugate formation relative to other events during intrinsic apoptosis. In RAW 264.7, HT-1080, and A549 cells, ABT/S6-induced conjugate formation occurred early and preceded CASP3 and PARP cleavage (**Fig. 3D,G; Fig. S3S**). Moreover, the pan-caspase inhibitor zVAD did not prevent conjugate formation (**Fig. 3E-G; Fig. S3D,E,I,J**). Thus, formation of the MIA40–TRX1 conjugate is an early post-MOMP event that is independent of caspase activation.

### MOMP reduces and/or destabilizes several MIA40 substrates

During these apoptosis experiments, we also monitored unconjugated MIA40 and COA7, a MIA40 substrate whose abundance is particularly sensitive to MIA40 inactivation (**Fig. S2S; Fig. S3S**)^44,45^, by immunoblotting. Because these samples were labeled with NEM, which preserves cysteine oxidation states without appreciably altering protein mass, oxidized and reduced forms of MIA40 and COA7 could be distinguished by their differential migration on nonreducing gels (**Fig. S4S**). Intriguingly, after MIA40–TRX1 conjugate formation, the remaining unconjugated MIA40 and COA7 migrated more slowly on nonreducing gels, consistent with reduction of their structural disulfides (**Fig. 3D,G; Fig. S3D,H,I,J**, arrowheads). Both proteins subsequently decreased in abundance, with COA7 showing more pronounced loss (**Fig 3D,G**), consistent with reduction preceding degradation. Together, these observations indicate that MOMP promotes the reduction and destabilization of at least some MIA40 pathway proteins.

To define MOMP-induced changes in protein abundance more comprehensively, we performed whole-cell quantitative MS-based proteomics in RAW 264.7 cells treated with ABT/S6. Remarkably, this treatment depleted only a small number of proteins, including CASP3, MIA40, AK2, and TXNIP (**Fig. 4A**, left). COA7 was not detected in the proteomics dataset, indicating that some MOMP-sensitive proteins were not captured by this analysis. We confirmed the loss of these proteins and COA7 by immunoblotting (**Fig. 4B**). The proteasome inhibitor bortezomib partially rescued their depletion (**Fig. 4A**, middle, **Fig. 4B; Fig. S4S**). Because bortezomib toxicity limited the concentration that could be used, these experiments likely underestimate the proteasomal contribution to this loss. Nevertheless, these data indicate that MOMP promotes proteasome-dependent degradation of select proteins.

**Figure 4.**
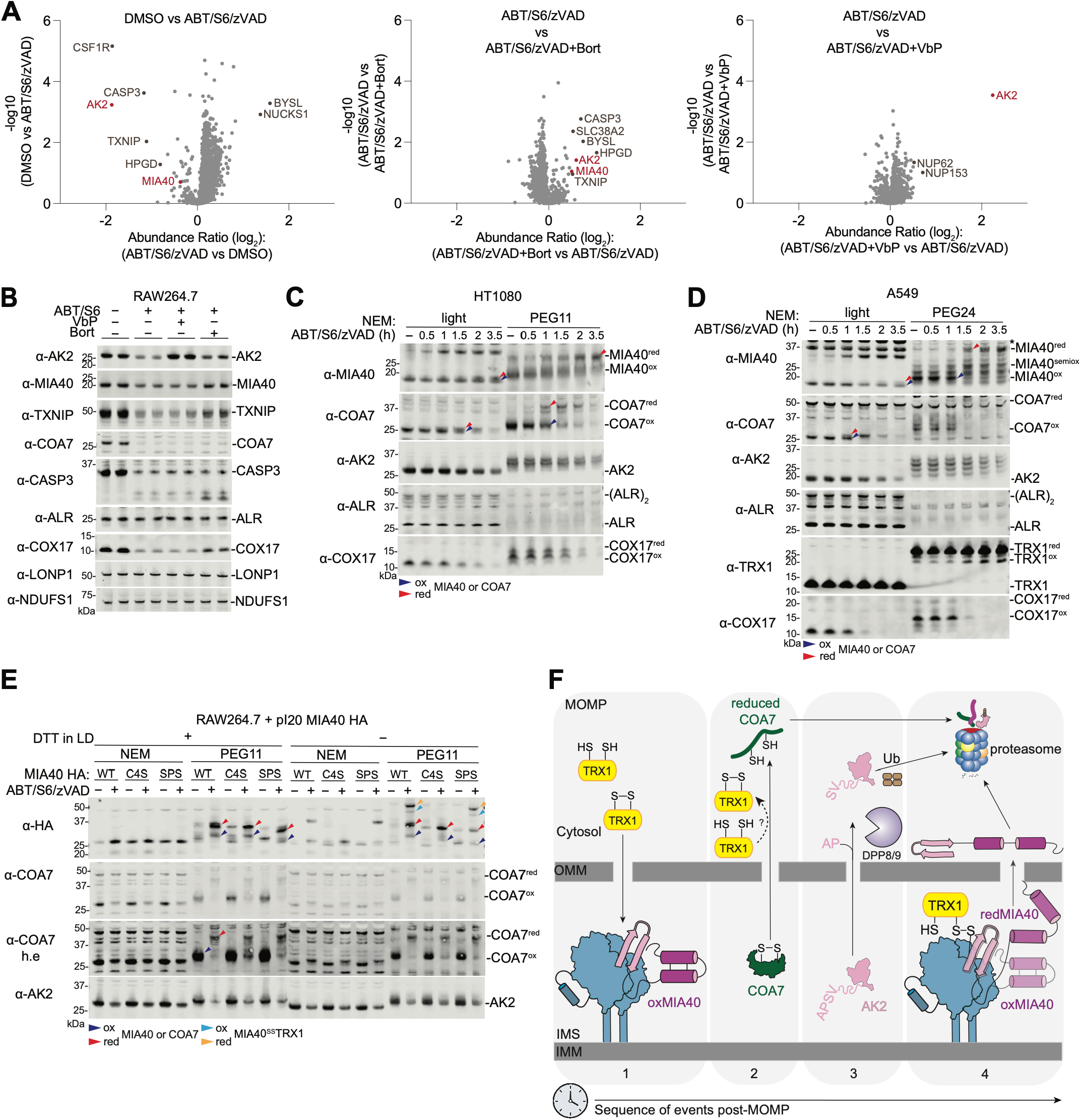
MOMP reduces and/or destabilizes several MIA40 pathway proteins. (**A–B**) Quantitative proteomic analysis of RAW264.7 cells following 3 h treatment with ABT/S6/zVAD alone (left), with bortezomib (middle), or with VbP (right) (**A**), with immunoblot analysis of selected proteins from the same samples (**B**). (**C–E**) Redox-state analysis following ABT/S6/zVAD treatment in HT-1080 cells (**C**), A549 cells (**D**), or RAW264.7 cells stably expressing the indicated MIA40 constructs (**E**); treatments were for the indicated times in (**C–D**) and for 3 h in (**E**). Arrowheads indicate the oxidized and reduced forms of MIA40 and COA7 and the corresponding MIA40–TRX1 conjugates, as indicated. ABT-263 (10 μM), S63845 (1 μM), zVAD (50 μM), VbP (10 μM), and bortezomib (500 nM) were used as indicated. Unless otherwise indicated, cells were lysed in buffer containing 40 mM NEM and samples were analyzed by immunoblotting under reducing or nonreducing conditions. For the PEG-based analyses in (**C–E**), samples were labeled with 15 mM PEG11 (**C, E**) or PEG24 (**D**). Samples in (**C–D**) were analyzed under reducing conditions, whereas samples in (**E**) were analyzed under both reducing and non-reducing conditions. (**F**) Proposed model for the sequential redox and proteostatic remodeling of the MIA40 pathway following MOMP. Immunoblots are representative of three or more independent experiments.

We next examined the cysteine oxidation states of COA7, MIA40, and AK2 by differential labeling of reduced cysteines with NEM or PEG-maleimide. Whereas NEM preserves oxidation-dependent migration differences on nonreducing gels, PEG-maleimide adds substantial mass to each reduced cysteine and therefore resolves reduced species on either reducing or nonreducing SDS–PAGE (**Fig. S4S**). As above, NEM-labeled COA7 and MIA40 migrated slightly more slowly on nonreducing gels following MOMP in multiple cell types, consistent with reduction of their structural disulfides (**Fig. 4C,D; Fig. S4C-E**, arrowheads). COA7 reduction occurred after approximately 1 h and was followed by rapid degradation, consistent with previous work showing that COA7 structural disulfides are critical for preventing misfolding and proteasome-mediated degradation^44^. MIA40 reduction occurred substantially later and its degradation was much less extensive. PEG-maleimide labeling more clearly resolved these changes as increased apparent molecular weight (**Fig. 4C-E; Fig. S4S**). Moreover, WT and Cys53Ser/Cys55Ser MIA40, but not Cys4Ser MIA40, showed substantial fractions in TRX1-conjugated and reduced states (**Fig. 4E; Fig. S4 C-E**, cyan and purple arrowheads), showing that the intermolecular TRX1–MIA40 disulfide remains stable even as the structural disulfides within MIA40 are reduced.

By contrast, AK2 showed little reduction following MOMP (**Fig. 4C-E; Fig. S4S**), perhaps because its structural disulfide is buried within the folded protein ^25^. Nevertheless, AK2 was robustly degraded, consistently after COA7 but before MIA40 (**Fig. 4A** left, B-E; **Fig. S4B-F**). We therefore asked whether DPP8/9 activity contributes to its degradation after MOMP^25^. Strikingly, the DPP8/9 inhibitor VbP selectively and nearly completely restored AK2 abundance following ABT/S6 treatment in RAW 264.7 (including NLRP1B-deficient), HT-1080, and A549 cells (**Fig. 4A** right, **4B; Fig. S4B-D,F**), demonstrating that AK2 stabilization is independent of inflammasome activation and occurs across multiple cell types. The structurally and mechanistically distinct DPP8/9 inhibitors 1G244, 8j, and CQ31 similarly stabilized AK2 during ABT/S6 treatment (**Fig. S4S**)^19,46^, as did bestatin methyl ester (MeBs), an inhibitor of the N-degron pathway^47^; by contrast, the control prolyl endopeptidase (PREP) inhibitor S17092 and the lysosomal inhibitor bafilomycin A1 had no effect on AK2 loss (**Fig. S4S**). Consistent with these data, DPP8/9 remained active during apoptosis, as measured using a fluorogenic GP-AMC reporter substrate (**Fig. S4S**). Together, these findings show that the DPP8/9-dependent N-degron pathway promotes AK2 degradation following MOMP^26^.

Finally, we examined the oxidation states of additional MIA40 substrates and of TRX1 by immunoblotting. ALR remained oxidized and stable following MOMP, whereas COX17 retained its disulfides but nevertheless underwent proteasome-dependent degradation (**Fig. 4B–D; Fig. S4S**). MIA40 substrates therefore differ substantially in their susceptibility to the redox and proteostatic changes that follow MOMP. A small fraction of TRX1 was oxidized in unstressed cells, and this fraction increased modestly after MOMP (**Fig. 4D, Fig. S4S**), indicating that TRX1 oxidation accompanies MIA40 substrate reduction. Overall, MOMP remodels the MIA40 oxidative-folding pathway through several distinct, largely sequential events: (1) oxTRX1 first conjugates to the oxidized MIA40–AIFM1 complex; (2) COA7 and potentially other IMS proteins undergo reduction and degradation; (3) AK2 is degraded through the DPP8/9-dependent N-degron pathway without detectable disulfide reduction; and (4) a subset of the MIA40 pool eventually undergoes partial structural disulfide reduction and degradation. Notably, however, a substantial fraction of MIA40 — with either intact or reduced structural disulfides — remains conjugated to TRX1 within the AIFM1 complex throughout (**Fig. 4F**).

### MIA40 oxidation and oxTRX1 binding promote AIFM1 dimerization

We next asked how MIA40 and TRX1 influence AIFM1 complex assembly. The prevailing model proposes that binding of NADH—and, less efficiently, NADPH^30^— induces AIFM1 dimerization, thereby exposing a surface that binds MIA40 or AK2 (**Fig. 1B**)^30–32^. We first tested this model by SEC. Purified AIFM1 eluted predominantly as a monomer and shifted substantially, but incompletely, toward the dimer upon the addition of stoichiometric NADH (**Fig. 5A; Fig. S5S**). Surprisingly, FL MIA40 alone shifted a fraction of AIFM1 into a MIA40-associated dimeric species in the absence of NADH (**Fig. 5A**). FL MIA40 and NADH together shifted AIFM1 almost completely into a dimeric, MIA40-bound species (**Fig. 5A**). WT MIA40 peptides spanning residues 2–17 or 2–30 bound AIFM1 but did not promote dimerization in the absence of NADH (**Fig. S5S**), indicating that regions outside the N-terminal β-hairpin are required for MIA40 alone to drive NADH-independent dimerization. Thus, MIA40 can bind AIFM1 in the absence of nucleotide, and FL MIA40 protein and NADH can cooperate to promote dimerization rather than acting through a strictly sequential assembly mechanism.

**Figure 5.**
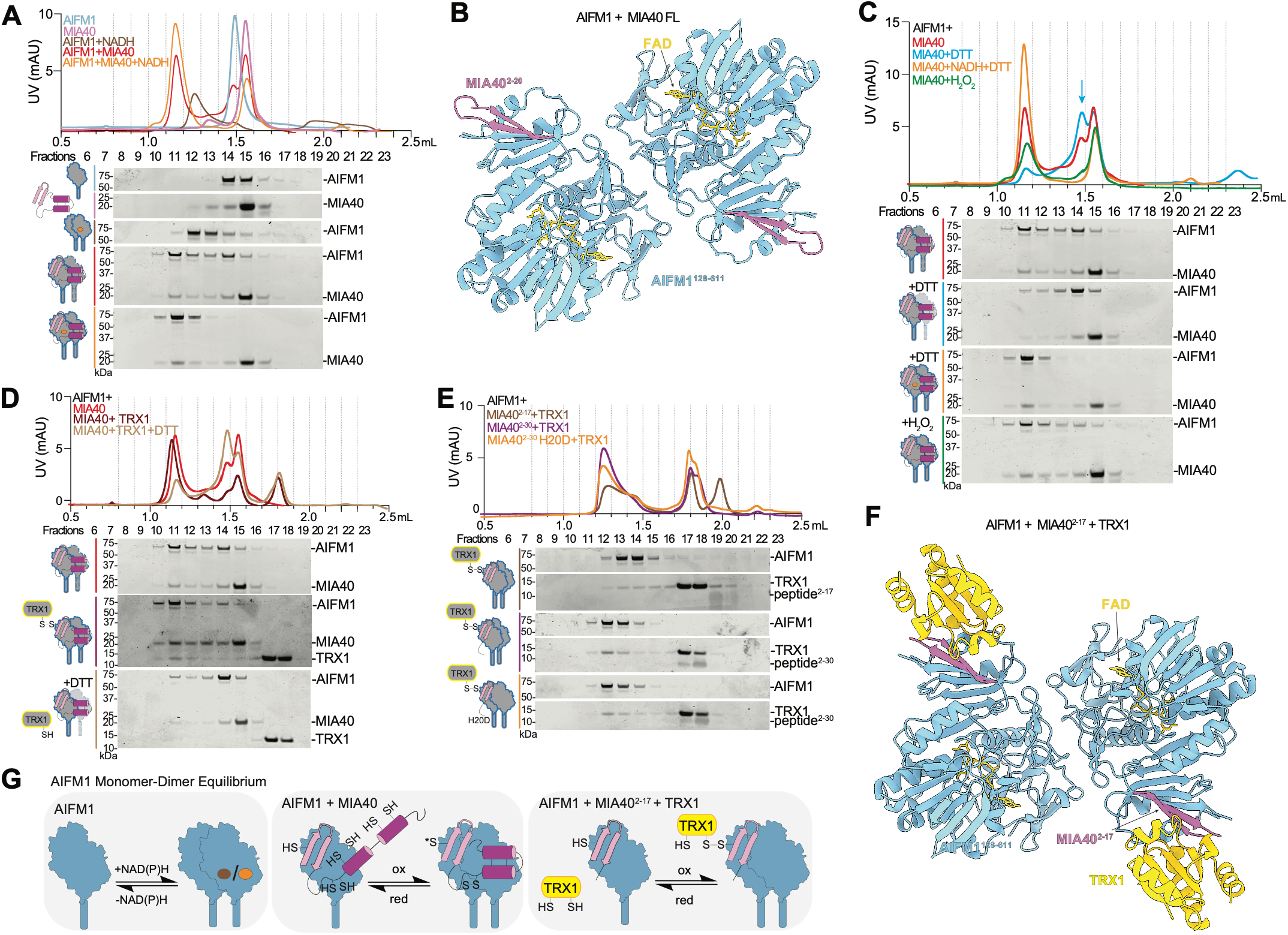
MIA40 oxidation and oxTRX1 binding promote AIFM1 dimerization. (**A**,**C-E**) SEC of the indicated proteins under the indicated conditions, Collected fractions were analyzed by Coomassie staining. The same AIFM1–MIA40 chromatogram/blot is shown in **A, C**, and **D**. Arrow in (**C**) indicates the MIA40-bound AIFM1 monomer. (**B**,**F**) Structure of the AIFM1–MIA40 Cys53Ser/Cys55Ser complex (PDB 38EB) (**B**) and the AIFM1– MIA40 peptide^2–17^–TRX1 complex (PDB 38ED) (**F**). (**G**) Proposed model for the AIFM1 monomer-dimer equilibrium. Asterisk denotes an oxidized thiol. Recombinant proteins were mixed at an AIFM1:MIA40:TRX1:peptide 1:2:4:10 molar ratio, with AIFM1 at 10 μM. NADH (10 μM), DTT (1 mM), and H_2_O_2_ (100 μM) were used. Chromatograms are representative of three or more independent experiments.

To characterize this NADH-independent complex structurally, we isolated an AIFM1 complex containing Cys53Ser/Cys55Ser MIA40, which yielded a more homogeneous SEC profile than WT MIA40 (**Fig. S5S**), and analyzed it by cryo-EM. Two particle populations were resolved in an approximately 6:1 ratio: a predominant AIFM1 dimer (**Fig. 5B; Fig. S5S** left) and a less abundant Ω-shaped dimer of dimers (**Fig. S5S** right, **Fig. S5S**). The predominant class yielded a 2.1 Å reconstruction with the canonical AIFM1 dimer architecture (**Fig. 5B**). Importantly, no NAD(H) density was detected in the nucleotide-binding site (**Fig. S5S**). The MIA40 N-terminal hairpin was resolved at the AIFM1 interface, but no additional MIA40 density was observed, leaving the structural basis by which regions outside the hairpin promote dimerization unresolved.

We reasoned that this NADH-independent dimer might not have been observed previously because earlier studies included TCEP^32^ or DTT^48^, whereas our experiments lacked reducing agents. Indeed, DTT attenuated formation of the NADH-independent MIA40–AIFM1 dimer and increased the later-eluting MIA40–AIFM1 monomer (**Fig. 5C**, arrow), whereas H_2_O_2_ favored dimer formation (**Fig. 5C**). PEG-maleimide labeling confirmed the redox sensitivity of Cys4 and the Cys53/Cys55 CPC motif and revealed minor species containing four or five reduced cysteines after DTT treatment, consistent with partial reduction of one or more structural disulfides (**Fig. S5S**). Notably, Cys4Ser MIA40 promoted substantially more AIFM1 dimerization than WT or Cys53Ser/Cys55Ser MIA40 under untreated, reducing, and oxidizing conditions (**Fig. S5S–J**), showing that the side chain at position 4 strongly influences the dimerization equilibrium. Nevertheless, WT, Cys4Ser, Cys53Ser/Cys55Ser, and Cys4Ser/Cys53Ser/Cys55Ser MIA40 proteins all promoted AIFM1 dimerization and remained at least partially responsive to DTT and H_2_O_2_ (**Fig. S5H–J**). Together, three observations suggest that one or more structural disulfides within the MIA40 substrate-binding domain contribute to the redox-dependent regulation of AIFM1 dimerization: redox sensitivity persisted in the triple mutant; DTT treatment generated species containing four or five reduced cysteines, including in the Cys53Ser/Cys55Ser mutant; and the N-terminal peptides failed to promote dimerization. Mutants that directly disrupted these structural disulfides could not be purified as homogeneous proteins, precluding direct evaluation of their contribution.

We next asked how TRX1 engagement of Cys4 affects AIFM1 dimerization. AIFM1 and TRX1 did not interact detectably in the absence of MIA40 (**Fig. S5S**). However, TRX1 bound the NADH-independent FL MIA40–AIFM1 complex and shifted AIFM1 further toward the dimeric state; this effect was reversed by DTT (**Fig. 5D**). Consistently, purified AIFM1, MIA40, and TRX1 formed a DTT-sensitive covalent ternary complex in the absence of NADH (**Fig. S1S; Fig. S2S**). Strikingly, co-incubation with TRX1 enabled the MIA40 N-terminal peptides—which did not promote AIFM1 dimerization on their own (**Fig. S5S**)—to form dimeric AIFM1 complexes (**Fig. 5E**). Cys4Ser peptides did not form these complexes (**Fig. S5S**). Even the weakly binding H20D MIA40 mutant peptide^32^ promoted AIFM1 dimerization in the presence of TRX1 (**Fig. 5E**), underscoring the potent dimerization-promoting effect of TRX1 engagement.

A 3.1 Å cryo-EM reconstruction of the AIFM1–MIA40 (2–17)–oxTRX1 complex revealed a symmetric AIFM1 dimer in which MIA40 Cys4 formed a disulfide bond with TRX1 Cys32, again without density attributable to NAD(H) (**Fig. 5F; Fig. S5S**). Notably, 3D classification did not identify a population of AIFM1 dimers lacking TRX1. Thus, ox-TRX1 engagement of MIA40 Cys4 is sufficient to enable the isolated MIA40 N-terminal β-hairpin alone to promote NADH-independent AIFM1 dimerization, bypassing the requirement for additional regions of MIA40. Together, these findings show that MIA40 and dinucleotide ligands can bind AIFM1 independently and cooperate to promote dimerization. They further identify two oxidation-dependent mechanisms that enhance MIA40-driven AIFM1 dimerization: oxidation of FL MIA40 and oxTRX1 engagement of MIA40 Cys4 (**Fig. 5G**).

### oxTRX1 binding shifts AIFM1 toward MIA40 over AK2

We next asked how oxTRX1 affects assembly of the competing AK2–AIFM1 complex. In the absence of dinucleotide, AK2 neither detectably associated with AIFM1 nor promoted its dimerization under basal, oxidizing, or reducing conditions (**Fig. S6S**). Upon addition of NADH, however, AK2 associated with AIFM1, and nearly all AIFM1 shifted into a dimeric, AK2-bound state (**Fig. S6S**). Although this result was consistent with cooperation between the two ligands, stoichiometric NADH alone shifted most AIFM1 into the dimeric state, leaving limited dynamic range for directly assessing cooperativity (**Fig. 5A**). We therefore turned to NADPH. At a stoichiometric concentration within the estimated cellular range^49^ (10 µM), NADPH alone did not detectably promote AIFM1 dimerization, and a 10-fold excess produced only a partial shift (**Fig. S6S**). In the presence of either AK2 or a MIA40 N-terminal peptide, however, 10 µM NADPH readily promoted dimerization (**Fig. S6S**), even though neither protein ligand promoted dimerization on its own. Thus, both AK2 and MIA40 N-terminal peptides cooperate with pyridine dinucleotides to stabilize dimeric AIFM1.

These results suggested that AIFM1 protein ligands differ in dimer-stabilizing potency, particularly when dinucleotides are limiting. Our data support a hierarchy of dimerization potency: AK2 and MIA40 N-terminal peptides were relatively weak, oxidized FL MIA40 was stronger, and oxTRX1-engaged MIA40 was most effective. Accordingly, in protein mixtures lacking added dinucleotide, MIA40-bound AIFM1 dimers predominated, and oxTRX1 further favored these complexes (**Fig. S6S**). At 10 µM NADPH, AK2 robustly formed a dimeric AK2–AIFM1 complex in the presence of a WT MIA40 N-terminal peptide. Addition of TRX1 markedly shifted AIFM1 toward the MIA40 peptide-bound complex, whereas a Cys4Ser peptide failed to support this redistribution (**Fig. 6A**). Similarly, when FL MIA40 was used, AK2 and WT MIA40 bound AIFM1 comparably in the absence of TRX1, whereas oxTRX1 further favored MIA40 binding (**Fig. S6S**). Thus, MIA40–TRX1 conjugation preferentially stabilizes MIA40-bound AIFM1 at low-to-moderate dinucleotide concentrations, even when only the MIA40 N terminus is present. At higher concentrations of NADH or NADPH, however, mixed AK2- and MIA40-bound complexes persisted even in the presence of oxTRX1 (**Fig. S6S**), indicating that increasing dinucleotide concentrations diminish the competitive advantage conferred by the stronger MIA40 ligands.

**Figure 6.**
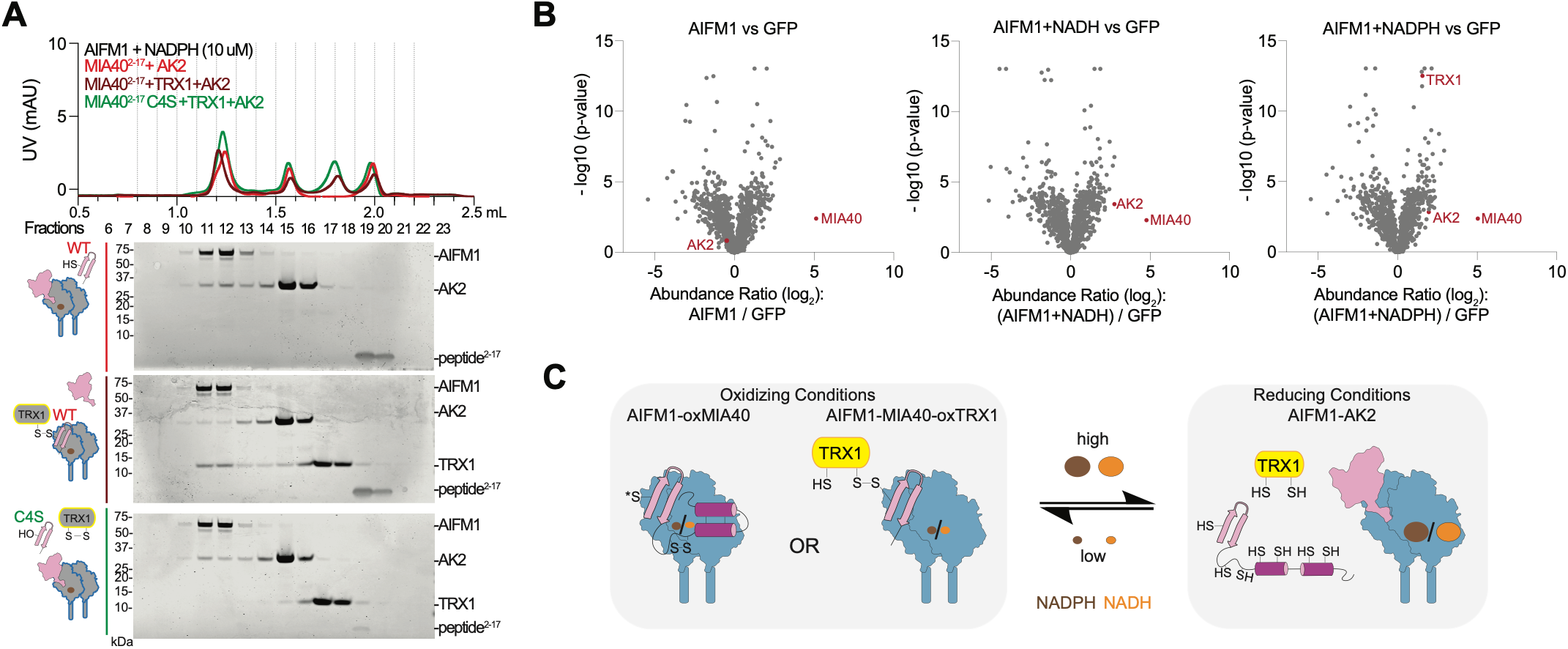
oxTRX1 binding shifts AIFM1 toward MIA40 over AK2. (**A**) SEC of recombinant AK2, AIFM1, TRX1, the indicated peptides, and NADPH as indicated. Collected fractions were analyzed by Coomassie staining. (**B**) Quantitative mass spectrometry of FLAG–AIFM1 immunoprecipitates in the absence or presence of NADH or NADPH (1 mM). (**C**) Proposed model for redox- and dinucleotide-dependent AIFM1 complex assembly. Oxidizing conditions favor MIA40-bound AIFM1, whereas reducing conditions favor AK2-bound AIFM1. Asterisk denotes an oxidized thiol. Recombinant proteins were mixed at an AIFM1:AK2:TRX1:peptide molar ratio of 1:2:4:10, with AIFM1 at 10 μM. Chromatograms are representative of three or more independent experiments.

Consistent with these biochemical data, FLAG-tagged AIFM1 immunoprecipitated MIA40 from HEK 293T cells without added dinucleotide, whereas AK2 association required addition of NADH or NADPH (**Fig. 6B; Fig. S6S**). These results suggest that the endogenous dinucleotide available in the lysates was insufficient to support robust AK2–AIFM1 binding. Together, these findings support a model in which oxidized MIA40 and limiting NAD(P)H favor MIA40-bound AIFM1, with oxTRX1 further stabilizing this state. Conversely, reduction of MIA40 and increasing NAD(P)H availability diminish this preference and permit AK2 association (**Fig. 6C**). Following MOMP, rapid AK2 degradation and TRX1–MIA40 conjugation would therefore act together to shift AIFM1 toward MIA40-bound states, particularly when NAD(P)H is limiting.

### Conjugate formation is insensitive to conventional redox perturbation

We tested whether redox-active compounds, respiratory inhibitors, or hypoxia altered MIA40–TRX1 conjugate formation following MOMP. None of these perturbations detectably altered ABT/S6-induced MIA40–TRX1 conjugate formation (**Fig. S7A–C**). H_2_O_2_ appeared to increase conjugate abundance, but also induced apoptosis on its own, suggesting that this effect may reflect increased MOMP rather than a direct increase in oxTRX1. Moreover, *TXNRD1*- knockout cells, which have elevated oxTRX1 levels (**Fig. S7S**), formed the conjugate following ABT/S6 treatment to a similar extent as WT cells (**Fig. S7S**), indicating that TRX1–MIA40–AIFM1 complex formation is not limited by the overall abundance of oxTRX1. DTT and NAC also failed to appreciably reduce the oxTRX1 pool (**Fig. S7S**), explaining the insensitivity of the TRX1–MIA40 conjugate to these treatments, and likewise did not alter inflammasome sensitivity to DPP8/9 inhibitors^10^. Collectively, these data suggest that the functionally relevant pool of oxTRX1 is buffered against conventional redox perturbations.

## Discussion

Here, we show that MOMP does more than release mitochondrial death factors: it initiates a coordinated redox and proteostasis remodeling program involving proteins that are normally segregated across the mitochondrial outer membrane (**Fig. 4F**). Following MOMP, cytosolic oxTRX1 immediately forms a stable intermolecular disulfide with MIA40 Cys4 at the MIA40–AIFM1 interface, promoting the formation of the MIA40-bound AIFM1 dimer. Subsequently, DPP8/9 cleave AK2 and target it for degradation, removing a competing AIFM1 ligand. Together, oxTRX1 and DPP8/9 stabilize the MIA40–AIFM1 complex. In parallel, at least one MIA40 substrate, COA7, undergoes reduction and degradation as TRX1 becomes progressively oxidized (**Fig. 4F**). At later times, MIA40 structural disulfides are partially reduced, yet the persistent Cys4–TRX1 disulfide indicates that a substantial fraction of MIA40 remains within the TRX1– MIA40–AIFM1 ternary complex.

The results reframe the DPP8/9–AK2 relationship. DPP8/9 were previously proposed to cleave AK2 that fails to be efficiently imported under basal conditions, but this quality-control pathway operates at relatively low flux, with only small amounts of cytosolic AK2 accumulating over days^25^. Following MOMP, by contrast, DPP8/9-dependent AK2 degradation is substantial, identifying apoptosis as a physiological context in which this pathway becomes strongly engaged. Intriguingly, DPP8/9 processing of human AK2 exposes an inhibitor of apoptosis protein (IAP) recognition site^26^, which could antagonize IAP function. Processed AK2 may therefore also contribute to apoptotic execution before its eventual proteasomal degradation.

Our results also refine the prevailing model of AIFM1 complex assembly. Previous work proposed that NADH binding first induces AIFM1 dimerization, thereby displacing the C-loop and exposing the shared MIA40- and AK2- binding interface (**Fig. 1B**) ^32,45^. We instead find that MIA40 can bind AIFM1 without dinucleotide, and that protein and dinucleotide ligands cooperate to promote dimerization, with members of each class differing markedly in potency. Among the protein ligands, oxidized FL MIA40 promotes AIFM1 dimerization more effectively than an isolated MIA40 N-terminal peptide or AK2. Engagement of MIA40 Cys4 by oxTRX1 provides a second, potent oxidation-dependent route to MIA40-driven AIFM1 dimerization (**Fig. 5G**). Among the dinucleotide ligands, NADH is substantially more potent than NADPH in driving dimerization^30^, but both can cooperate with protein ligands to stabilize dimeric AIFM1. Thus, AIFM1 complex assembly is not a strictly dinucleotide-first pathway, but instead reflects a tunable equilibrium jointly determined by the relative MIA40 and AK2 availability, MIA40 redox state, oxTRX1 engagement, and dinucleotide levels (**Fig. 6C**).

Because association with dimeric AIFM1 enhances MIA40’s catalytic activity^32,34,50^, this assembly mechanism could tune MIA40-mediated oxidative folding during apoptosis. Indeed, substantial pools of folded MIA40 and ALR persist after MOMP, suggesting that the oxidative-folding machinery could retain at least partial functional capacity. We therefore propose the following model. First, MOMP immediately exposes pre-existing IMS protein disulfides to cytosolic TRX1, whose oxidation during substrate reduction could supplement the basal oxTRX1 pool^51^. oxTRX1 binding and AK2 degradation stabilize dimeric, MIA40-bound AIFM1 and could sustain MIA40-dependent oxidative folding after MOMP. This sustained activity could provide a second, continuing source of oxTRX1: MIA40 installs new substrate disulfides that TRX1 subsequently reduces. The resulting oxTRX1 could, in turn, stabilize MIA40–AIFM1, establishing a feed-forward cycle that sustains oxidative folding and maintains NLRP1 restraint during apoptosis (**Fig. 7A**, “feed-forward cycle”). This model predicts that disrupting the ternary complex would weaken this cycle, accelerating the reduction and degradation of MIA40 and its substrates, shifting TRX1 toward a more reduced state, and potentially facilitating NLRP1 activation. Consistent with part of this prediction, COA7 and MIA40 were reduced and degraded more rapidly following MOMP in AIFM1-deficient cells, which cannot form the ternary complex (**Fig. 3D**). AIFM1 loss alone, however, did not eliminate the oxTRX1 pool (**Fig. S7S**), impair apoptotic progression, or induce pyroptosis (**Fig. 3D**), indicating that additional coordinated changes are required to generate the proposed NLRP1-activating danger state.

**Figure 7.**
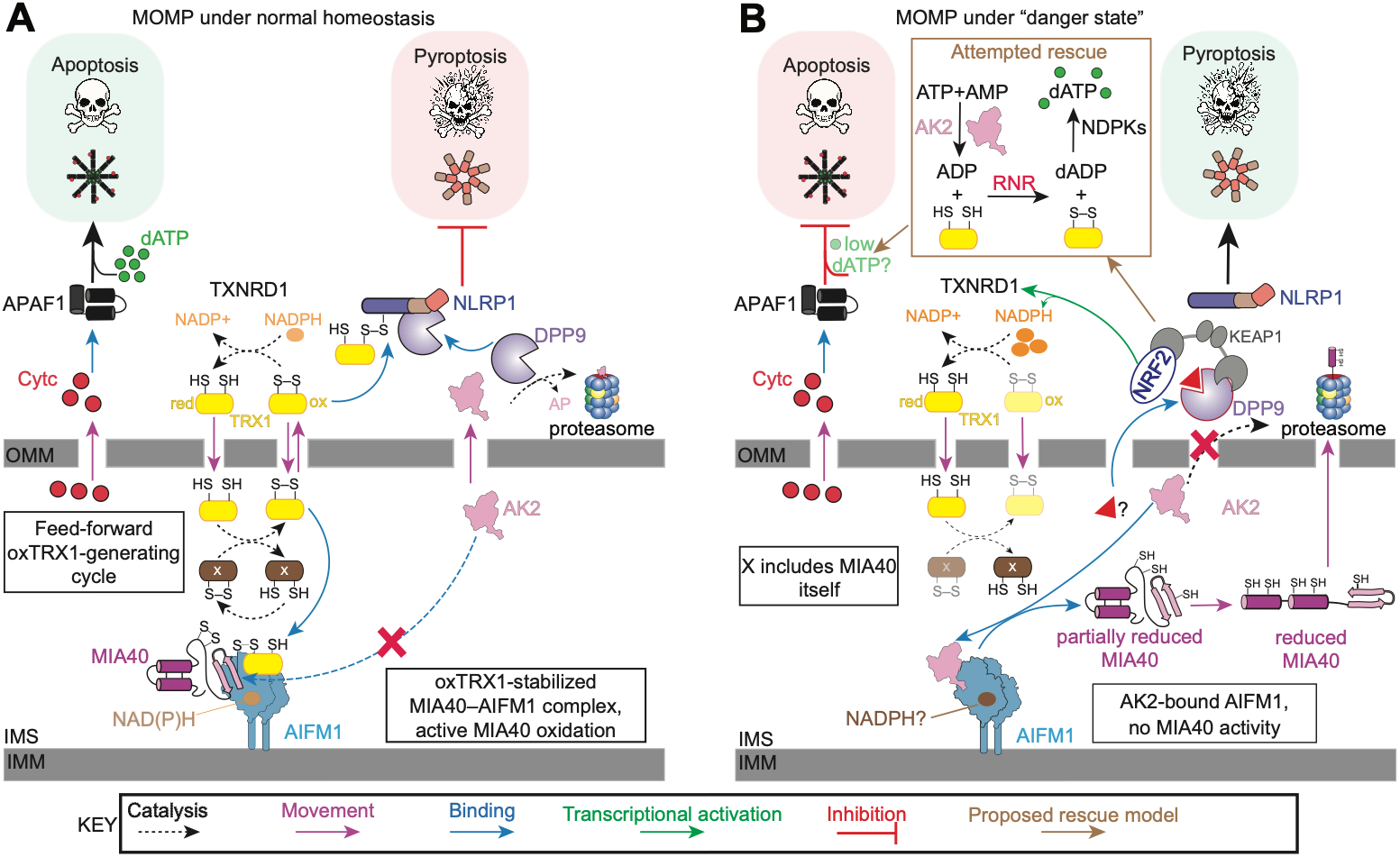
Proposed redox and proteostasis remodeling following MOMP. (**A**) During normal apoptotic execution, MOMP releases cytochrome c (Cytc) to drive APAF1-dependent apoptosis, while oxTRX1 and active DPP9 restrain NLRP1. These same factors stabilize the MIA40–AIFM1 complex: oxTRX1 binds the complex directly, whereas DPP8/9 activity removes the competing ligand AK2, together helping maintain a relatively oxidizing state via a possible “feed-forward” cycle. (**B**) In the proposed “danger state” in which apoptotic execution is impaired, engagement of the DPP9–KEAP1 complex stabilizes AK2 and activates NRF2-dependent reducing programs, increasing redTRX1 and favoring AK2-bound AIFM1. We speculate that stabilized AK2 and redTRX1 may support RNR-dependent dATP production, potentially compensating for dATP levels insufficient for apoptosome activation. These same changes would weaken oxTRX1- and DPP9-dependent NLRP1 restraint, potentially triggering inflammasome activation if apoptotic competence is not restored.

What physiological condition might generate this state? The post-MOMP timing of the DPP8/9-dependent remodeling described here suggests a possible connection to impaired apoptotic execution (**Fig. 7B**). Apoptosome activation requires cytochrome c together with dATP or, at substantially higher concentrations, ATP^52,53^, and several factors implicated here intersect with deoxynucleotide biosynthesis. AK2 contributes to the ADP pool that ribonucleotide reductase (RNR) converts to dADP, a reaction supported by reducing equivalents supplied through TRX1 and ultimately NADPH ^54^. Consistent with the importance of this axis for nucleotide supply, loss of either AK2 or TXNRD1 causes severe hematopoietic proliferative defects, with both perturbations disrupting nucleotide homeostasis and TXNRD1 deficiency directly limiting RNR-dependent dNTP synthesis ^55,56^. Thus, if dATP availability limits efficient apoptosome activation, mobilizing AK2 and redTRX1 could represent an attempted rescue mechanism (**Fig. 7B**).

One route to mobilizing these factors would be engagement of the mutually inhibitory DPP9–KEAP1 complex^21^. DPP9 inhibition would stabilize AK2 and prolong its enzymatic activity, while KEAP1 inhibition would activate NRF2 and increase reducing capacity, including TXNRD1 expression and NADPH availability^57,58^. Increased NADPH, reduced TRX1, and stabilized AK2 could also shift AIFM1 away from MIA40 and weaken the proposed feed-forward oxTRX1 cycle (**Fig. 7A**). These changes could therefore favor dATP synthesis while simultaneously eroding both established restraints on NLRP1. In this model, NLRP1 would respond to a persistent, unsuccessful attempt to restore apoptotic competence by redirecting the cell toward pyroptosis.

More broadly, our study reveals a coordinated signaling program downstream of MOMP that operates independently of apoptotic caspase activation. Strikingly, several of the same proteins that restrain inflammatory cell death also participate in the remodeling events accompanying intrinsic apoptosis. By exposing pre-existing protein disulfides and favoring MIA40–AIFM1 assembly, MOMP could help maintain NLRP1 restraint and thereby reinforce the typically noninflammatory mode apoptotic death, at least under normal conditions. Understanding these post-MOMP changes and the conditions that disrupt them may help explain why vertebrate intrinsic apoptosis is organized around mitochondrial permeabilization—an arrangement that is not universal across metazoans and may therefore represent a later layer of metabolic regulation imposed on a more ancient caspase-based death program^59^. Such an arrangement could couple the threshold, kinetics, and mode of cell death to metabolic state, favoring immunologically silent apoptosis when execution proceeds efficiently and inflammatory pyroptosis when it does not.

### Limitations of the study

Several mechanistic and physiological aspects of this work remain unresolved. First, although we showed that MIA40- driven AIFM1 dimerization is redox dependent, we could not determine whether or how the MIA40 substrate-binding domain mediates this activity. This region was not resolved in our cryo-EM reconstruction, and mutations directly disrupting its structural disulfides caused MIA40 to aggregate during purification. Further biochemical and structural studies will therefore be required to define this mechanism. Second, although we establish that the TRX1–MIA40– AIFM1 ternary complex forms immediately after MOMP, we do not yet know how TRX1 engagement affects MIA40 oxidative-folding activity or AIFM1 NADH oxidase activity^34^ in the post-MOMP cellular environment. Our data also do not establish whether TRX1 is principally responsible for reducing MIA40 substrate proteins after MOMP or exclude contributions from other reducing systems, including glutathione. Third, we relied on *AIFM1* knockout as a proxy for loss of the TRX1–MIA40–AIFM1 complex after MOMP. Although it eliminates this ternary complex, this perturbation does not recreate the other proposed features of the NLRP1-activating danger state, such as engagement of the DPP9–KEAP1 complex. These changes may need to occur together to fully reduce TRX1 and to redirect cell death responses. Defining the physiological conditions that generate this danger state— and determining whether nucleotide insufficiency is one such trigger—will require further investigation.

## Acknowledgements

This work was supported by the NIH (R01 AI137168, R01 AI163170, R01 CA266478 to D.A.B.; the MSKCC Core Grant P30 CA008748; F99 CA305522 to L.P.T), Marie-Josée Kravis Women in Science (WISE) fellowship (L.P.T.), The Starr Foundation Basic Research Innovation Award Grant (M.B.G.), and Mr. William H. and Mrs. Alice Goodwin and the Commonwealth Foundation for Cancer Research and The Center for Experimental Therapeutics of Memorial Sloan Kettering Cancer Center (D.A.B.). We thank the Patel laboratory for providing access to equipment, Dr. Richard Hite for helpful cryo-EM advice, and Dr. Mara Monetti and Zhuoning Li for MS assistance. This paper was typeset with the bioRxiv word template by @Chrelli: www.github.com/chrelli/bioRxiv-word-template.

## Author contributions

D.A.B. conceived and directed the project. L.P.T., M.B.G, M.T., Z.L., A.T., and D.P.B. performed cloning, gene editing, biochemistry, and cell biology experiments, designed experiments and analyzed data. L.P.T, M.T., B.L., A.C., and S.S. performed cryo-EM experiments. B.L., S.S., and M.J.d.l.C collected and analyzed cryo-EM data. L.P.T. and D.A.B. wrote the manuscript. All authors reviewed and provided input to the manuscript.

## Materials and Methods

### Antibodies and reagents

The primary antibodies that were used in this study are the following: anti- CHCHD4 polyclonal antibody (MIA40; Proteintech, 21090-1-AP), anti- CHCHD4 monoclonal antibody (Proteintech, 66718-1-Ig), anti-AIF rabbit monoclonal antibody (Cell Signaling Technology, 5318), anti-FLAG antibody (Millipore Sigma, F1804), anti-HA antibody (Cell Signaling Technology, C29F4), anti-V5 antibody (Sigma-Aldrich, V8012), anti-Thioredoxin 1 antibody (Cell Signaling Technology, 2429S) for human TRX1 and anti-TRX/Thioredoxin antibody (Abcam, ab273877) for mouse TRX1, anti-GAPDH antibody (Cell Signaling Technology, 2118S), anti-β-actin antibody (Sigma-Aldrich, A5441), anti-PARP antibody (Cell Signaling Technology, 9542S), anti-CASP3 antibody (Cell Signaling Technology, 9662S), anti-cleaved CASP3 antibody (Cell Signaling Technology, 9661S), anti-BAX antibody (Proteintech, 50599-2-Ig), anti-BAK antibody (Proteintech, 29552-1- AP), anti-cytochrome c antibody (Proteintech, 66264-1-Ig), anti-NLRP1 antibody (R&D Systems, AF6788), anti-GSDMD antibody (Abcam, ab209845), anti-COA7 antibody (Proteintech, 25361-1-AP), anti-AK2 antibody (Proteintech, 11014-1-AP), anti-TXNIP antibody (Cell Signaling Technology, 14715S), anti-ALR/GFER antibody (Proteintech, 11293-1-AP), anti-TRIAP1 antibody (Proteintech, 15351-1-AP), anti-COX17 antibody (Proteintech, 11464-1-AP), anti-DPP9 antibody (Abcam, ab42080), anti-NDUFS1 antibody (Proteintech, 12444-1-AP), and anti-LONP1 antibody (Proteintech, 15440-1-AP). The secondary antibodies that were used in this study are the following: IRDye 800CW donkey anti-rabbit (LI-COR, 925-32211), IRDye 680RD donkey anti-rabbit (LI-COR, 925-68073), IRDye 800CW donkey antimouse (LI-COR, 925-32212), and IRDye 680RD donkey anti-mouse (LI-COR, 925-68072) secondary antibodies were used for immunoblotting. Anti-FLAG M2 affinity resin (Millipore Sigma, A2220), anti-FLAG M2 magnetic beads (Sigma-Aldrich, M8823), and Pierce IP Lysis Buffer (Thermo Scientific, 87787) were used for FLAG immunoprecipitation. N-ethylmaleimide (NEM; Sigma-Aldrich, E3876-5G), DL-dithiothreitol (DTT; Sigma-Aldrich, D9779), tris(2-carboxyethyl)phosphine hydrochloride (TCEP; Sigma-Aldrich, C4706), iodoacetamide (IAA; Millipore Sigma, I6125), PEG11-maleimide (Thermo Scientific, 21911), PEG24-maleimide (Thermo Scientific, 22713), PEG24-maleimide (BroadPharm, BP-22750), NADH (Fisher Scientific, AAJ6163803), and NADPH (Millipore Sigma, 481973-100MG) were used as indicated. Val-boroPro (VbP; Tocris, 3719), bestatin methyl ester (MeBs; Sigma, 200485), 1G244 (Fisher Scientific, 50-258-1317), 8j, CQ31 (synthesized in-house), S17092 (Sigma-Aldrich, SML0181-5MG), bortezomib (Millipore Sigma, 504314), bafilomycin A1 (Cayman Chemical Company, 11038), subasumstat (MedChemExpress, HY-111789), zVAD-fmk (MedChemExpress, HY-16658B-10MG), and VX-765 (Fisher Scientific, 50- 101-3602), staurosporine (Selleck Chemicals, S1421), etoposide (Fisher Scientific, AAJ63651MB), raptinal (MedChemExpress, HY-121320-10MG), ABT-737 (ApexBio, A8193), ABT-263 (Selleck Chemicals, S1001), S63845 (Selleck Chemicals, S8383), dexamethasone (Sigma-Aldrich, D4902- 25MG), LPS (InvivoGen, TLRL-3PELPS), nigericin sodium salt (Cayman Chemical Company, 11437), imiquimod (Cayman Chemical Company, 14956), RSL3 (Cayman Chemical Company, 19288), H_2_O_2_ (Sigma-Aldrich, 95321), and hydroxyurea (HU; MedChemExpress, HY-B0313) were used as indicated. GP-AMC (Cayman Chemical Company, 34458) and zGP-AMC (MedChemExpress, HY-D1670) were used for DPP8/9 and PREP activity assays, respectively. Dynabeads Human T-Activator CD3/CD28 for T Cell Expansion and Activation (Fisher Scientific, 11-131-D), recombinant human IL-2 (PeproTech, 200-02), CytoTox-Fluor Cytotoxicity Assay (Promega, G9262), and Mitochondria Isolation Kit for Cultured Cells (Thermo Scientific, 89874) were used as indicated. Doxycycline hyclate (Cayman, 14422), FuGENE HD (Promega, E2311), and HALT protease inhibitor cocktail (Thermo Scientific, 78430) were used according to the manufacturers’ instructions. 4X Protein Sample Loading Buffer for Western Blots (LI-COR Biosciences, 928-40004), NuPAGE Sample Reducing Agent (10X; Thermo Fisher Scientific, NP0009), NativePAGE Sample Buffer (4X; Thermo Fisher Scientific), NativePAGE™ G-250 Sample Additive (Thermo Fisher Scientific), and Intercept (PBS or TBS) Blocking Buffer (LI-COR) were used for SDS-PAGE, BN-PAGE, and immunoblotting as indicated. InstantBlue Coomassie Protein Stain (Abcam, ab119211) and Protein Standard Mix, 15–600 kDa, for size-exclusion chromatography (Sigma-Aldrich, 69385-30MG), were used for gel staining and SEC analysis. Triethylammonium bicarbonate (TEAB; Thermo Scientific, 90114), HEPES Buffer (1 M, pH 8.0; Boston BioProducts, BBH-80), Trypsin/Lys-C Mix, Mass Spec Grade (Promega, V5071), TMT10plex Isobaric Label Reagents and Kits (Thermo Fisher Scientific, 90110), Pierce™ C18 Tips (Thermo Scientific, 87784), Pierce Peptide Desalting Spin Columns (Thermo Scientific, 89852), and Pierce High pH Reversed-Phase Peptide Fractionation Kit (Thermo Scientific, 84868) were used for peptide cleanup and fractionation prior to mass spectrometry. SHuffle T7 Express E. coli cells (New England Biolabs), Rosetta™2(DE3) competent cells (Novagen/EMD Millipore, 71397-3), isopropyl-β-D-thiogalactopyranoside (IPTG; GoldBio, I2481C5), Pierce Protease Inhibitor Tablets, EDTA-free (Thermo Scientific, A32965), HisPur Ni-NTA Resin (Thermo Scientific, 88223), and ULP1 protease (produced in-house), were used for bacterial protein expression and purification.

### Cell culture

HEK293T, RAW264.7, HT-1080, and A549 cells were purchased from ATCC and cultured in Dulbecco’s Modified Eagle’s Medium (DMEM) with L-glutamine supplemented with 10% fetal bovine serum (FBS). N/TERT-1 keratinocytes, a gift from James Rheinwald, were cultured in Keratinocyte Serum-Free Medium (KSFM; Gibco) supplemented with 100 U/mL penicillin, 100 µg/mL streptomycin, bovine pituitary extract (20 µg/mL; Gibco), and recombinant epidermal growth factor (EGF; 2 ng/mL; Gibco), or, when unavailable, in Defined Keratinocyte SFM (Gibco) with the same supplements. Frozen human peripheral blood mononuclear cells (PBMCs) were purchased from STEMCELL Technologies (70025) and cultured in Roswell Park Memorial Institute 1640 medium (RPMI-1640) supplemented with 10% FBS. Cryopreserved purified human CD3 pan T cells were purchased from IQ Biosciences (IQB-HU1- T50). T cells were thawed in RPMI-1640 medium supplemented with 10% FBS and 30 U/mL IL-2 (PeproTech). T cells were activated for 48 h using Human T-Activator CD3/CD28 Dynabeads for T-Cell Expansion and Activation (Gibco) according to the manufacturer’s protocol. All cells were maintained at 37 °C in a humidified atmosphere containing 5% CO_2_. Cell lines were regularly tested for mycoplasma contamination using the MycoAlert Mycoplasma Detection Kit (Lonza).

### Generation of knockout cell lines

CRISPR-based genome editing was used to generate the knockout cell lines used in this study. MIA40 (CHCHD4)-deficient HT-1080 cells and BAX/BAK double-knockout RAW264.7 cells were generated using Cas12a with the all-in-one pRB054 vector containing the indicated guide RNAs. AIFM1- and TXNRD1-deficient RAW264.7 cells were generated using ribonucleoprotein (RNP)-based genome editing. NLRP1B-deficient RAW264.7 cells were generated using Cas9. Edited cells were expanded as single-cell clones or bulk populations and knockout was confirmed by immunoblotting.

### Cloning and site-directed mutagenesis

The MIA40 open reading frame (ORF) was purchased from OriGene (NM_001098502, RC217831), and the AIFM1 ORF (pENTR-AIF) was purchased from Addgene (plasmid #16182). The TRX1 ORF was obtained from Dan Ball et al. For mammalian expression, TRX1, MIA40 (CHCHD4), and AIFM1 constructs were subcloned into pDONR221 and transferred by Gateway cloning (Thermo Fisher Scientific) into pLEX307-derived vectors containing FLAG, Myc, HA, or V5 epitope tags, or into pInducer20-HA (Addgene #44012) for doxycycline-inducible expression. For bacterial expression, constructs were generated in pRSF-SUMO-Duet vectors using restriction enzyme cloning. Site-directed mutagenesis was performed using the QuikChange II Site-Directed Mutagenesis Kit (Agilent, 200523) according to the manufacturer’s instructions for both mammalian and bacterial expression constructs. Mutants used in this study included TRX1 C32S, C35S, C32S/C35S, and C62S/C69S/C73S, as well as MIA40 C4S, C53S, C55S, C53S/C55S (SPS), C4S/C53S/C55S, and CS7. The AIFM1–MIA40 fusion construct used for structural studies was generated as previously described _32_.

### Stable cell line generation and transient transfection

Expression constructs were packaged into lentivirus in HEK293T cells using psPAX2 (Addgene #12260) and pMD2.G (Addgene #12259). Viral supernatants were collected 48 h after transfection and added to the indicated target cells. Polybrene was included at 1 µg/mL for all cell lines except RAW264.7 cells. One day after transduction, cells were replated and antibiotic selection was initiated. Cells were selected with puromycin at 0.5 µg/mL for HT-1080 cells or 5 µg/mL for RAW264.7 cells, or with G418 at 400 µg/mL, as appropriate, until control cells were eliminated. For transient transfections, HEK293T or HT-1080 cells were plated either in 12-well plates at 0.25 × 106 cells/mL or in 6-well plates at 0.5 × 106 cells per well and transfected the following day with the indicated plasmids (1 or 2 µg total DNA per well), using pLEX307-RFP as filler DNA where necessary to maintain equal total DNA amounts. Transfections were performed using FuGENE HD (Promega) according to the manufacturer’s instructions, and cells were harvested 48 h after transfection. For experiments using doxycycline-inducible constructs, expression of the indicated constructs was induced with doxycycline (100 ng/mL) for 24 h before treatment.

### Immunoprecipitation

FLAG immunoprecipitations were performed using anti-FLAG M2 magnetic beads (Sigma-Aldrich, M8823), except for the experiment shown in **Figure 1F**, for which anti-FLAG M2 affinity resin (Millipore Sigma, A2220) was used. Unless otherwise indicated, cells expressing the indicated FLAG-tagged constructs were harvested and lysed by sonication. For the experiments shown in **Figures 6B** and S6F, cells were instead lysed without sonication in Pierce™ IP Lysis Buffer (Thermo Scientific, 87787) supplemented with Halt Protease, Pierce Universal Nuclease (1:1,000), and 1 mM NADH or NADPH, as indicated, for 15 min on ice with periodic vortexing. Clarified lysates were incubated with FLAG beads either for 1 h at room temperature or overnight at 4 °C. Beads were washed three times with PBS or TBS containing 0.1% Tween-20 (TBST), and bound proteins were eluted with 3×FLAG peptide (100 ng/µL) for 1-2 h at room temperature before downstream immunoblot or mass spectrometry analysis. For the on-bead reduction experiment shown in **Figure S1S**, beads were first washed and then incubated with 50 mM DTT for 20 min at room temperature prior to elution.

### Mitochondrial fractionation

Mitochondrial and cytosolic fractions were prepared from cultured cells using the Mitochondria Isolation Kit for Cultured Cells (Thermo Scientific, 89874) according to the manufacturer’s instructions, using the reagent-based differential centrifugation method. NEM was added, and fractions were subsequently analyzed by immunoblotting under the indicated reducing or nonreducing conditions.

### Solubility assay

Cells were lysed as indicated, and whole-cell lysates were centrifuged at 21,000 × g for 30 min at 4 °C to separate soluble and insoluble material. The supernatant was collected as the soluble fraction, whereas the pellet was retained as the insoluble fraction. The insoluble pellet was resuspended in the same volume represented in the soluble fraction, and equal volumes of each fraction were analyzed by SDS-PAGE and immunoblotting.

### TMT mass spectrometry of immunoprecipitated proteins

FLAG-Immunoprecipitated proteins were eluted from beads in 50 mM HEPES buffer (pH 8.5) containing 150 ng/µL 3XFLAG peptide (Sigma-Aldrich, F4799). Urea and TCEP were added to final concentrations of 2 M and 6 mM, respectively, and samples were shaken at 1,300 rpm for 15 min. Iodoacetamide was then added to a final concentration of approximately 18 mM, and samples were shaken at 1,300 rpm for 30 min in the dark. Proteins were digested with Trypsin/Lys-C Mix (2 µg) for 3 h at 37 °C. Samples were diluted with 50 mM EPPS (pH 8.5) to reduce the urea concentration to <1 M and incubated overnight at 37 °C with shaking at 1,300 rpm. Digested peptides were desalted using Pierce™ C18 Tips (Thermo Fisher Scientific, 87784) or Pierce™ Peptide Desalting Spin Columns (Thermo Fisher Scientific, 89852) according to the manufacturer’s instructions. Desalted peptides were resuspended in 50 mM EPPS (pH 8.5) and labeled with TMT10plex™ Isobaric Label Reagents (0.2 mg per sample) in acetonitrile for 1 h at room temperature. Labeling reactions were quenched with hydroxylamine at a final concentration of 0.5% and incubated for 15 min at room temperature. For the experiments shown in **Figures 1C** and **1G**, samples were pooled and fractionated using the Pierce™ High pH Reversed-Phase Peptide Fractionation Kit (Thermo Fisher Scientific, 84868). Labelled peptides were resuspended in 0.1% formic acid and 1 µg was analyzed by mass spectrometry.

### Mass spectrometry of whole cell lysate

RAW264.7 cells were rinsed with PBS, detached from the plate by scraping, and lysed by sonication. Lysates were clarified by centrifugation at 1,000 × g for 5 min, and protein amounts were normalized to 100 µg per sample. Samples were adjusted to 50 mM TEAB and 2 M urea, reduced with 10 mM TCEP, and shaken at 1,300 rpm for 15 min. Iodoacetamide was then added to a final concentration of 20 mM, and samples were shaken at 1,300 rpm for 30 min in the dark. Proteins were digested with Trypsin/Lys-C Mix (2 µg) for 3 h at 37 °C. Samples were diluted with 50 mM TEAB to reduce the urea concentration to <1 M and incubated overnight at 37 °C with shaking at 1,300 rpm. Digested peptides were desalted using Pierce™ C18 Tips (Thermo Fisher Scientific, 87784) according to the manufacturer’s instructions. Desalted peptides were dried under vacuum and resuspended in 0.1% formic acid. Peptide concentration was measured using a NanoDrop spectrophotometer, samples were normalized, and 1 µg of peptide was analyzed by mass spectrometry.

### Mass spectrometry data analysis

Thermofisher RAW files were analyzed using MaxQuant (V2.1.0.0) or proteome discover.

### Immunoblotting and Blue Native-PAGE (BNP)

Protein concentrations were determined and normalized using the DC Protein Assay (Bio-Rad). Unless otherwise indicated, lysis was performed in the presence of 40 mM NEM or 15 mM PEG-maleimide, as indicated in the figure legend of each experiment. For SDS-PAGE, a 2X loading dye was prepared by combining 4X Protein Sample Loading Buffer for Western Blots (LI-COR Biosciences, 928-40004), NuPAGE Sample Reducing Agent (10X; Thermo Fisher Scientific, NP0009), and water at a 3:2:5 ratio. Samples were mixed 1:1 with the 2X loading dye before electrophoresis. For non-reducing conditions, an equivalent volume of water was substituted for the Nu- PAGE™ Sample Reducing Agent. Samples were separated on 4–12% Bis-Tris SDS-PAGE gels (Thermo Fisher Scientific), and proteins were transferred to nitrocellulose membranes using the Trans-Blot Turbo Transfer System (Bio-Rad) for 7 min at 2.5 A, up to 25 V. Blue Native-PAGE (BN-PAGE) was performed as previously described21. Briefly, FLAG-purified protein samples were combined 1:1 with a master mix containing 4X NativePAGE™ Sample Buffer, H_2_O, and NativePAGE™ G-250 Sample Additive (Thermo Fisher Scientific), to a final G-250 concentration of 0.25%. Samples were separated on 4–16% NativePAGE™ gels, first using dark blue cathode buffer containing 0.02% G-250 for 30 min at 150 V, followed by light blue cathode buffer containing 0.002% G-250 for 30 min at 300 V. Proteins were transferred to methanol-activated PVDF membranes using the Trans-Blot Turbo Transfer System for 10 min at 2.5 A, up to 25 V. Membranes were destained in 50% methanol and 10% acetic acid in water. For both SDS-PAGE and BN-PAGE immunoblotting, membranes were blocked with Intercept™ (TBS) Blocking Buffer (LI-COR) for 30 min at room temperature and incubated with primary antibodies overnight at 4 °C. Membranes were washed with TBS containing 0.1% Tween-20, incubated with fluorophore-conjugated secondary antibodies for 1 h at room temperature, washed again, and imaged using an Odyssey Imaging System (LI-COR).

### AMC-substrate assays

For lysate (WCL) AMC reporter assays, approximately 15 µg of total lysate was used and 100 µM GP-AMC or zGP-AMC. Assays were performed 384- well, black, clear bottom plates (Corning). Fluorescence was recorded using either Cytation five Cell Imaging Multi-Mode Reader 500 (BioTek) or Synergy H1 Multimode Microplate Reader (BioTek).

### Purification of recombinant proteins

The N-terminal His-tagged TRX1 expression plasmid pET16b-Trx was obtained from Addgene (plasmid #36944) and used as the source of the TRX1 ORF. Full-length MIA40, AK2, AIFM1(104–613), TRX1(C62S/C69S/C73S), and AIFM1(118–613) fused to MIA40(1–45) through an SGSGPGSGS linker were codon-optimized for E. coli expression and synthesized by GenScript. All recombinant protein constructs were cloned into pRSF-Duet-His_6_- SUMO, a modified pRSF-Duet-1 vector encoding an N-terminal His_6_- SUMO tag cleavable by ubiquitin-like protease 1 (ULP1). Plasmids encoding TRX1, AK2, and MIA40 constructs were transformed into E. coli SHuffle T7 Express cells (New England Biolabs), whereas plasmids encoding AIFM1 constructs were transformed into E. coli Rosetta 2(DE3) cells (Novagen). Single colonies were used to inoculate overnight starter cultures, which were subsequently expanded in LB medium containing kanamycin (50 µg/mL). Cultures were grown at 37 °C to an OD600 of approximately 0.6. For AIFM1 expression, LB medium was supplemented with 20 µM riboflavin and 25 µM FAD. Protein expression was induced with 0.5 mM IPTG, and cultures were incubated for 16 h at 16 °C with shaking. Cells were harvested by centrifugation at 3,500 g for 10 min and lysed using an EmulsiFlex-C3 high-pressure homogenizer (Avestin) in buffer containing 25 mM Tris-HCl (pH 8.0), 500 mM NaCl, 5% glycerol, 20 mM imidazole, 1 mM TCEP, and protease inhibitors. Lysates were clarified by centrifugation at 24,000 × g for 30 min (twice), and His_6_-SUMO-tagged proteins were purified by immobilized metal affinity chromatography using Ni-NTA resin at 4 °C in the presence of 1 mM TCEP. Bound proteins were eluted in lysis buffer containing 250 mM imidazole and 1 mM TCEP. The His_6_-SUMO tag was removed by incubation with ULP1 protease in the presence of 1 mM TCEP, and the cleavage reaction was passed over Ni-NTA resin a second time, also in the presence of 1 mM TCEP, to remove the His_6_-SUMO tag, His-tagged ULP1, and uncleaved protein. Purified proteins were subsequently subjected to size-exclusion chromatography using a Superdex 200 10/300 GL column (Cytiva) equilibrated in 25 mM Tris-HCl (pH 7.7) and 150 mM NaCl, either in the absence or presence of 1 mM DTT.

### Size exclusion chromatography (SEC)

Analytical SEC experiments were performed on an ÄKTA Pure system using a Superdex 200 Increase 3.2/300 GL column (Cytiva) equilibrated in 25 mM Tris-HCl (pH 7.5) and 150 mM NaCl. Recombinant proteins were combined at the indicated molar ratios using 5X stock solutions and incubated for 15 min at room temperature before analysis. Where indicated, DTT (1 mM), H_2_O_2_, (100 µM), NADH (10 or 100 µM as indicated), or NADPH (10 or 100 µM as indicated) were added directly to the protein mixture. DTT, NADH, and NADPH were included in the SEC running buffer and the column was equilibrated with the corresponding buffer for 1CV before sample injection. H_2_O_2_, was only added in the protein mixture and was not included in the running buffer. Samples (12 µL) were injected using a 50 µL sample loop and separated at 4 °C at a flow rate of 0.05 mL/min. Protein elution was monitored by absorbance at 280 nm. Fractions were collected in 100 µL volumes. For analysis of SEC fractions, samples were mixed 1:1 with loading dye, and 60 µL was loaded onto 4–12% NuPAGE™ Bis-Tris Midi Protein Gels, 1.0 mm, 20-well WedgeWell format.

### PEG-maleimide labelling of recombinant MIA40

Purified MIA40 WT, Cys4Ser, and Cys53Ser/Cys55Ser proteins were diluted to 100 µM working stocks and added to 15 µL reactions at a final concentration of 20 µM, matching the MIA40 concentration used in the SEC reconstitution experiments. Protein mixtures were treated with the indicated concentrations of H_2_O_2_ (10–100 µM) or DTT (1–10 mM) and then mixed 1:1 with PEG24-maleimide for a final concentration of 40 mM. An equal volume of loading dye was subsequently added, samples were boiled, resolved by SDS-PAGE and analyzed by Coomassie staining, with approximately 1.4 µg of MIA40 loaded per lane.

### Cryo-EM sample preparation and data collection

For the AIFM1^104-613^–MIA40^1–45^–TRX1 sample shown in **Figure 1H**, purified AIFM1–MIA40 fusion protein at 5 mg/mL was mixed with purified TRX1(C62S/C69S/C73S) at a 1:1.5 molar ratio in buffer containing 25 mM Tris (pH 7.7), 150 mM NaCl, and 1 mM NADH. Residual DTT from protein purification was present at approximately 1 mM in the final sample. Immediately before grid preparation, (1H,1H,2H,2H-perfluorooctyl)-β-D-maltopyranoside (FOM) was added to a final concentration of 0.5 mM. A 4 µL aliquot of the sample was applied to glow-discharged UltrAuFoil R1.2/1.3 300-mesh gold grids. Grids were blotted for 2 s at a blot force of 0, 6 °C, and 100% humidity and plunge-frozen in liquid ethane using a Vitrobot Mark IV (Thermo Fisher Scientific). A data set of 8,306 movies was collected on a Titan Krios G4 300 kV microscope equipped with a Falcon 4i camera and Selectris X energy filter (Thermo Fisher Scientific) using a 10-eV slit width, at the MSKCC Structural Biology Core (RRID: SCR 027809). Movies in EER format (2.93 seconds and 900 frames) were collected using EPU with a 0.725 Å pixel size, −0.5 to −1.7 µm defocus range, and 60 e^-^ Å^-^2 total dose per exposure. For the AIFM1^104-613^–MIA40 full-length complex shown in **Figure 5B**, AIFM1 and full-length MIA40 C53S/C55S were mixed at a 1:2 molar ratio, corresponding to approximately 250 µM AIFM1 and 500 µM MIA40. No DTT was present in this sample. The mixture was subjected to size-exclusion chromatography, and fraction 11 containing the AIFM1–MIA40 complex was collected for cryo-EM grid preparation. No FOM was added to this sample. A 4 µL aliquot was applied to glow-discharged UltrAuFoil R1.2/1.3 300-mesh gold grids, blotted for 2 s at a blot force of 0, 4 °C, and 100% humidity, and plunge-frozen in liquid ethane using a Vitrobot Mark IV. A data set of 7,289 movies was similarly collected except using a −0.5 to −1.8 µm defocus range. For the AIFM1^104-613^–MIA40^2–17^–TRX1 complex shown in **Figure 5F**, purified AIFM1, TRX1, and MIA40^2–17^ peptide were mixed at a 1:4:10 molar ratio to final concentrations of approximately 100 µM AIFM1, 400 µM TRX1, and 1 mM MIA40 peptide in a total volume of 20 µL. No DTT was present in this sample. FOM was added to a final concentration of 0.5 mM immediately prior to grid preparation. No size-exclusion chromatography was performed before grid freezing. A 3.5 µL aliquot of the sample was applied to glow-discharged UltrAuFoil R1.2/1.3 300-mesh gold grids. Grids were blotted for 2.5 s at a blot force of 0, 4 °C, and 100% humidity and plunge-frozen in liquid ethane using a Vitrobot Mark II. A data set of 5,364 movies was collected on a Titan Krios G2 300 kV microscope (Thermo Fisher Scientific) equipped with a Gatan K3 camera (Gatan). Movies (1.52 seconds and dose-fractionated into 38 frames) were collected using SerialEM ^60^ with a 0.642 Å pixel size and a −0.7 to −1.7 µm defocus range, with a total dose of and 55.3 e^-^ Å^-^2 per exposure.

### Cryo-EM data processing, model building, and refinement

All cryo-EM data processing was carried out with CryoSPARC (v4.7.1) ^61^. For the AIFM1^104-613^–MIA40^1–45^–TRX1 data set, movies were gain-corrected, fractionated into 45 EER fractions, frames aligned in Patch Motion Correction, and CTF parameters were estimated with Patch CTF Estimation. Particle autopicking was performed on denoised micrographs, and extracted particles underwent one round of 2D classification. Particles in 2D classes with clear structural features were selected for a four-class *ab-initio* reconstruction, producing one class resembling an AIFM1 dimer and three amorphous classes. These references were used for heterogeneous refinement, which was performed on all extracted particles except those corresponding to ice contaminants from 2D classification. Particles classifying to the AIFM1 dimer class were retained for a second round of *ab-initio* reconstruction and heterogeneous refinement. Because the TRX1 density was weaker than the adjacent AIFM1 density, the particles were C2 symmetry-aligned and refined by non-uniform refinement, then subjected to a four-class 3D classification; one class with approximately 25% of the particles contained strong TRX1 density on both sides of the dimer, while the other three classes lacked one or both copies of TRX1. The substoichiometry of TRX1 may be a result of residual reducing agent carried over from steps prior to sample preparation. Topaz picking ^62^ was trained on a subset (~8000) of the particles with full TRX1 occupancy, and the Topaz-extracted particles underwent the same classification procedure described. Particles from the classes containing fully-bound TRX1 of the two 3D classifications were combined and duplicate particles were removed. The resulting 304,502 particles were split into beam-image shift groups and refined by C2 symmetry non-uniform refinement with higher-order aberration correction and per-particle defocus optimization to yield a 2.8 Å reconstruction. The AIFM1^104-613^– MIA40 full-length data set was similarly processed, and 1,138,466 particles of the AIFM1^104-613^–MIA40 dimer were refined in C2 symmetry to 2.1 Å resolution. Due to anisotropy evident in the initial reconstruction of the AIFM1^104-613^–MIA40 tetramer, the Rebalance Orientations job was used to remove particles in over-represented orientation bins; the remaining 157,901 particles were refined in C2 symmetry to 3.0 Å resolution. The AIFM1^104-613^–MIA40^2–17^–TRX1 data set was similarly processed, with the C2 reconstruction from 437,255 particles extending to 3.2 Å resolution. Subsequent 3D classification of the data failed to identify particles with substoichiometric TRX1, even though the same procedure could resolve the compositional heterogeneity of TRX1 in the AIFM1^104-613^–MIA40^1–45^–TRX1 data. Final maps were generated from processing of the CryoSPARC output sharpened maps with EMReady ^63^. Initial models were built by docking the cryo-EM structure of the AIFM1–MIA40 complex (9GQZ) ^34^ and the crystal structure of TRX1 (1ERT) ^40^ into their respective maps with UCSF Chimera ^64^, with the exception of the AIFM1104-613–MIA402-17–TRX1 complex, which was built using the AIFM1104-613–MIA401-45–TRX1 complex from this study. Models underwent iterative rounds of manual adjustment in Coot ^65^, and real space refinement and MolProbity structure validation ^66^ in PHENIX ^67^.

### Statistical analysis

Two-sided Student’s t-tests were used for significance testing. *P* values less than 0.05 were considered to be significant. Graphs and error bars represent means ± SEM of three independent experiments unless stated otherwise. The investigators were not blinded in all experiments. All statistical analysis was performed using GraphPad Prism 9 or R.

### Declaration of generative AI and AI-assisted technologies in the manuscript preparation process

During the preparation of this work, the authors used Claude and ChatGPT for editing assistance. The authors reviewed and edited the output as needed and take full responsibility for the content of the published article.

---

**Figure S1.**
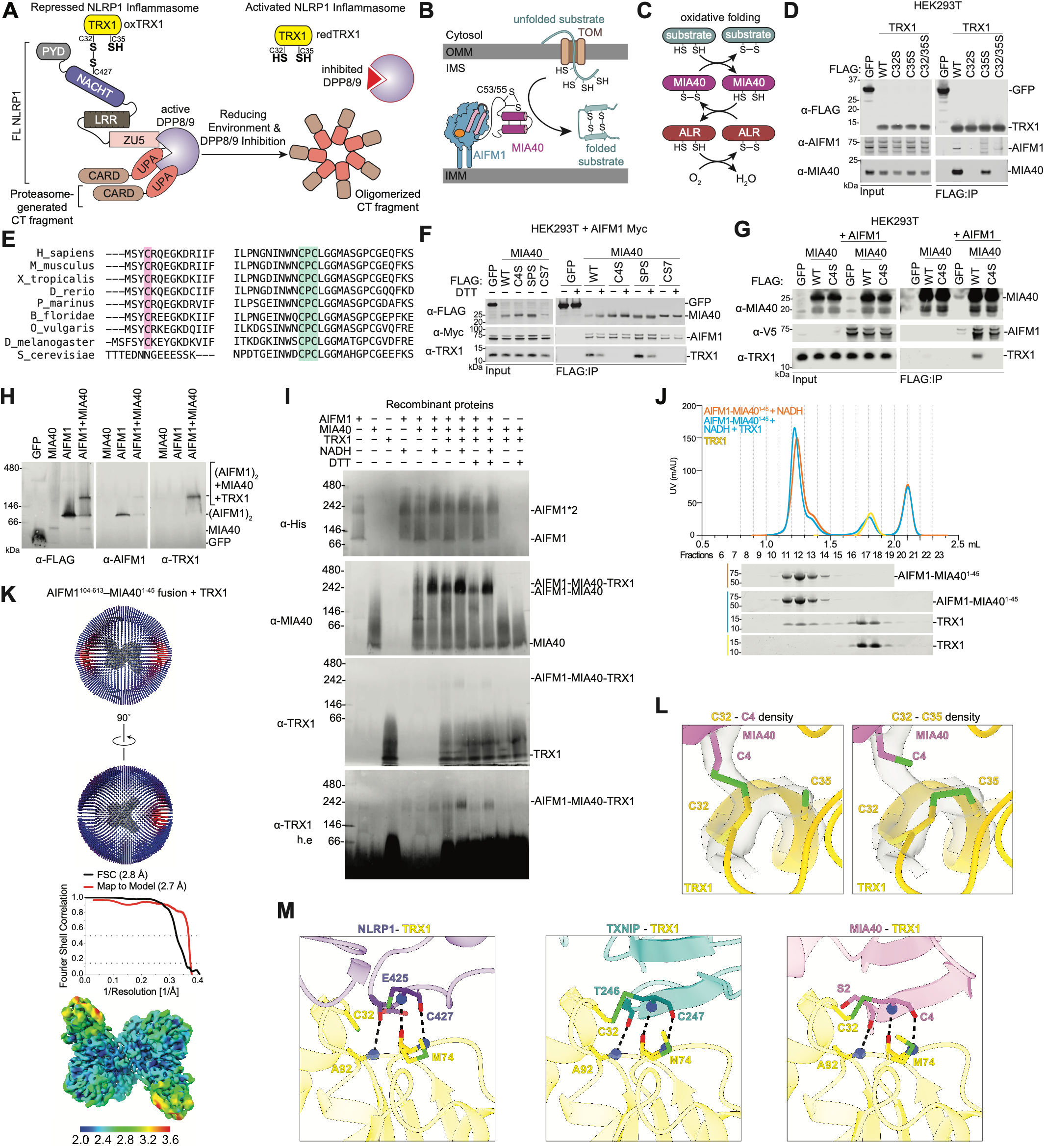
Characterization of the TRX1–AIFM1–MIA40 complex. (**A**) NLRP1 inflammasome regulation. NLRP1 autoproteolysis generates Nterminal (NT) and C-terminal (CT) fragments that remain noncovalently associated. oxTRX1 binding protects the NT fragment from degradation, whereas DPP8/9 bind FL NLRP1 together with the proteasome-released CT in a ternary complex that sequesters the CT. A reducing environment and DPP8/9 inhibition can relieve these restraints, leading to CT oligomerization and inflammasome activation. (**B**–**C**) MIA40-dependent mitochondrial protein import (**B**) and oxidative folding (**C**). (**D**,**F**) The indicated proteins were expressed in HEK293T cells, IPed, and analyzed by immunoblotting. In (**F**) the beads were treated with or without DTT (50 mM) before immunoblotting. (**E**) Sequence alignment of MIA40 residues surrounding C4 and the CPC motif across the indicated species. (**G**) Immunoblots corresponding to **Figure 1G** samples. (**H**,**I**) BNP immunoblot analyses of the indicated protein complexes isolated from HEK 293T cells (**H**) or assembled using recombinant AIFM1, MIA40, and TRX1 (Cys 62/69/73 mutated to Ser) ± NADH (100 μM) and DTT (10 mM) (**I**). (**J**) Size-exclusion chromatography of the AIFM1–MIA40^1–45^ fusion protein (50 μM) ± TRX1 (Cys62/69/73Ser, 120 μM) and NADH (100 μM). (**K**) Cryo-EM data processing and validation for the TRX1–MIA40–AIFM1 complex and local resolution estimate of the cryo-EM map. (**L**) Cryo-EM density corresponding to either an intermolecular MIA40 Cys4–TRX1 Cys32 disulfide (left) or an intramolecular TRX1 Cys32–Cys35 disulfide (right). (**M**) Structural comparison of the TRX1 interfaces with NLRP1, TXNIP, and MIA40 (dotted lines, hydrogen bonds), showing conserved backbone hydrogen bond interactions flanking their respective disulfide cysteine residues. Immunoblots and chromatograms are representative of three or more independent experiments.

**Figure S2.**
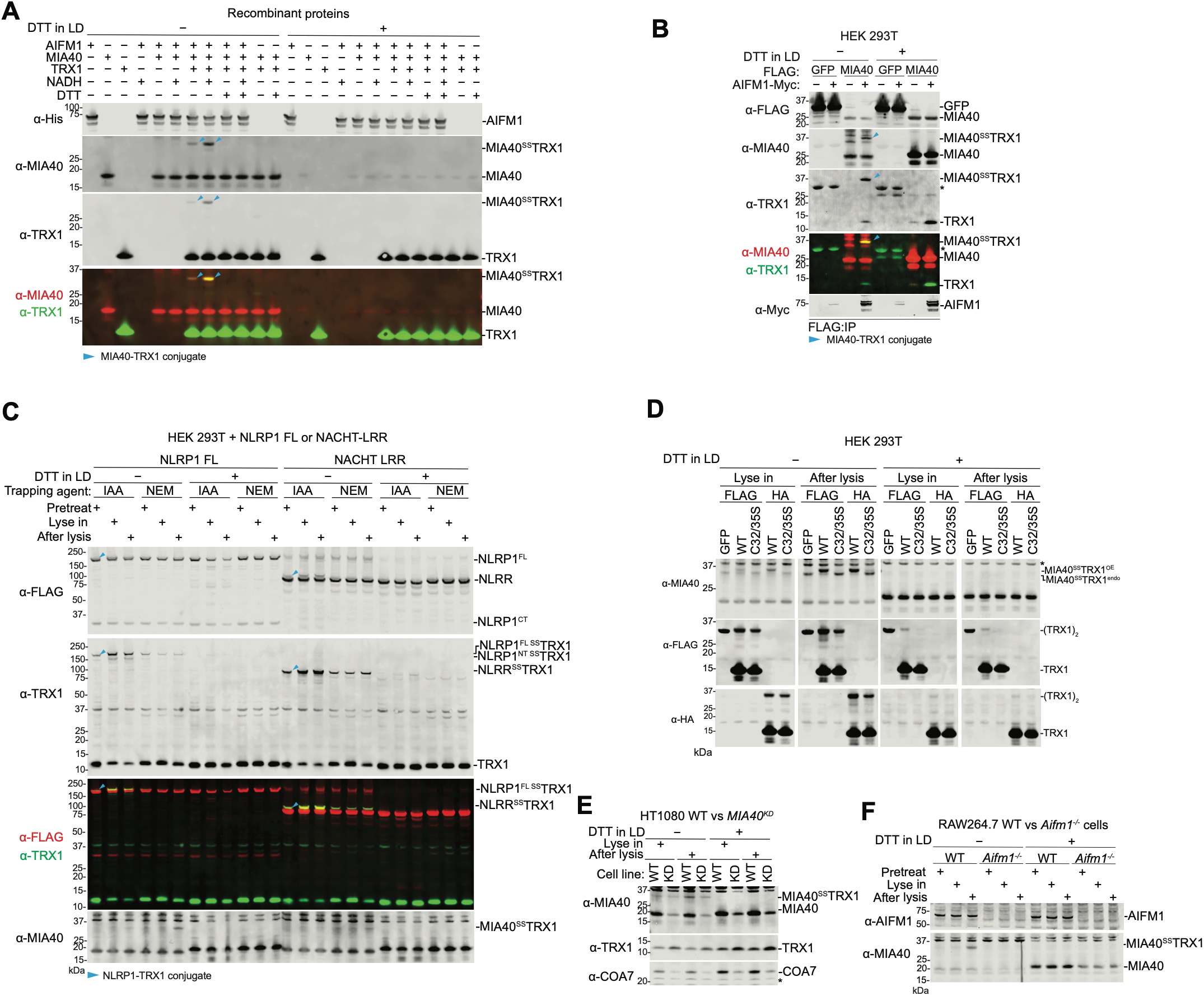
Characterization of the TRX1–MIA40 conjugate. (**A**) Recombinant AIFM1, MIA40, and TRX1 (Cys 62/69/73 mutated to Ser) were combined under the indicated DTT (10 mM) and NADH (100 μM) conditions, trapped with NEM (40 mM), and analyzed by immunoblotting under reducing or nonreducing conditions. Cyan arrowheads indicate MIA40–TRX1 conjugates. (**B–F**) FLAG-IPs from HEK 293T cells expressing the indicated proteins (**B**–**D**), WT and MIA40-deficient HT1080 cells (**E**), and WT and AIFM1-deficient RAW264.7 cells (**F**) were subjected to the indicated NEM (40 mM) and IAA (10 mM) thiol-trapping conditions and analyzed by immunoblotting under reducing or nonreducing conditions. Arrowheads indicate the MIA40–TRX1 or the NLRP1 FL– or NACHT LRR–TRX1 conjugates. Immunoblots are representative of three or more independent experiments. Asterisks denote non-specific bands.

**Figure S3.**
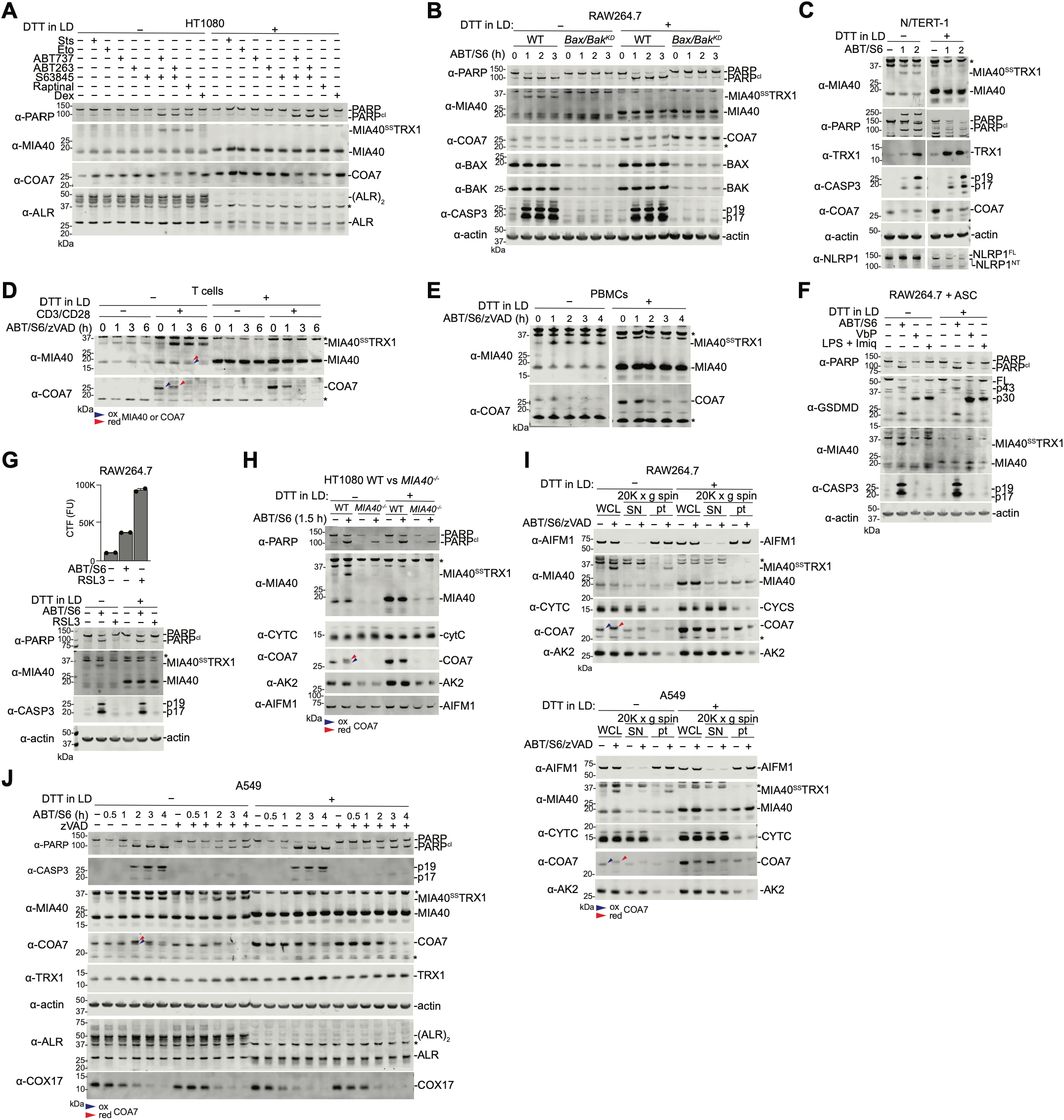
Intrinsic apoptosis induces MIA40-TRX1 conjugate formation. (**A**–**J**) The indicated cells were treated with the specified compounds for 2 h (**A**,**I**), the indicated intervals (**B-E, H**,**J**), or 3 h (**F**,**G**). All cells were lysed in buffer containing 40 mM NEM, except in **C** was in 10 mM IAA, and samples were analyzed by immunoblotting under reducing or nonreducing conditions. In (**G)**, cell death was additionally measured by CytoTox-Fluor fluorescence. In (**I**), lysates were separated into soluble and precipitate fractions by centrifugation. Compound concentrations: staurosporine (10 μM), etoposide (20 μM), ABT-737 (10 μM), ABT-263 (10 μM), S63845 (1 μM), raptinal (10 μM), dexamethasone (50 μM), zVAD (50 μM), VbP (10 μM), LPS (1 μg/mL), imiquimod (100 μM), and RSL3 (10 μM). Immunoblots are representative of three or more independent experiments. Asterisks denote non-specific bands.

**Figure S4.**
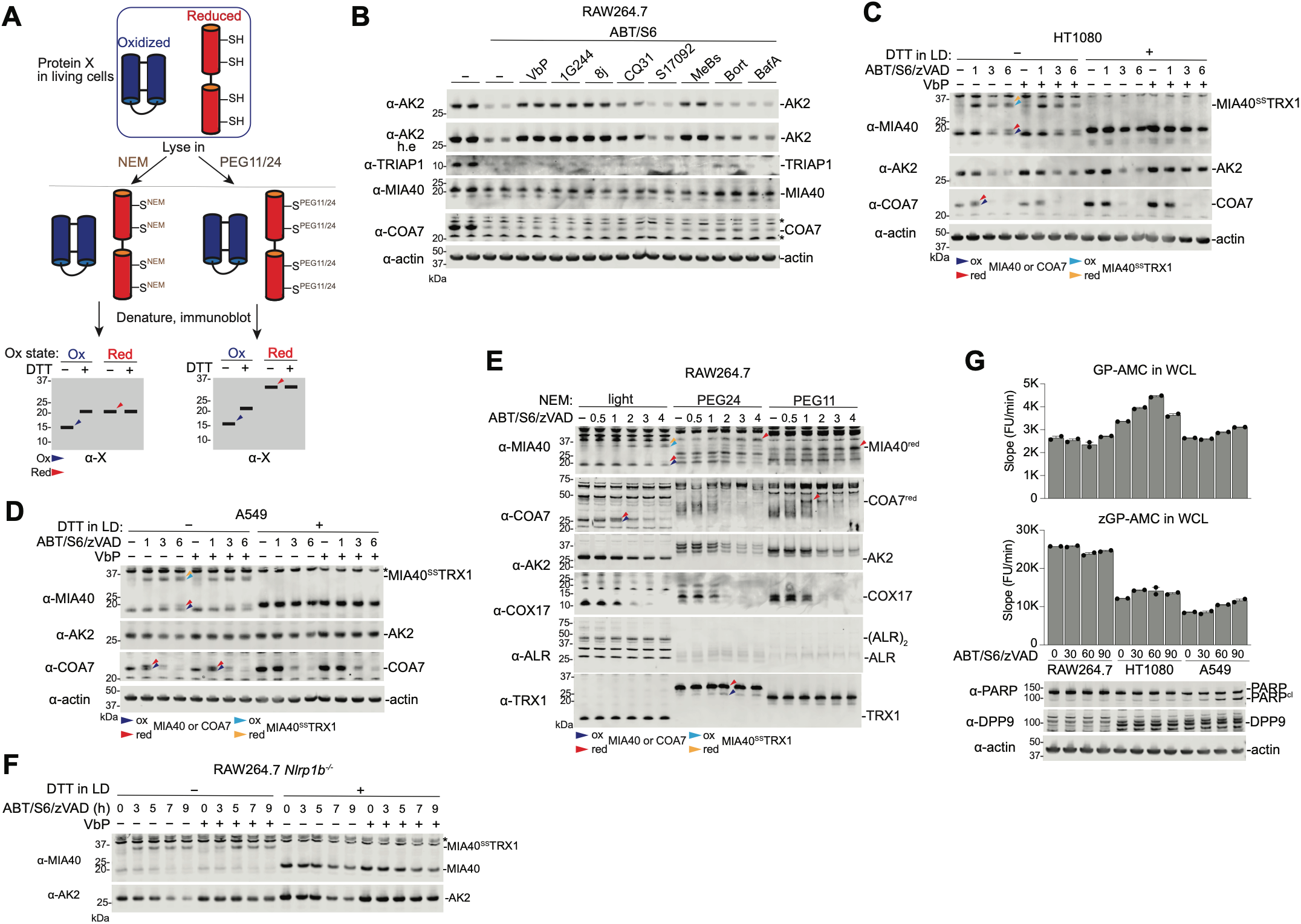
DPP8/9 inhibition blocks AK2 degradation after MOMP. (**A**) Schematic of NEM- and PEG-based thiol-labeling assays to distinguish oxidized and reduced protein species. (**B**) RAW264.7 cells were co-treated with zVAD, VX-765, and the indicated compounds for 3.5 h. (**C–F**) The indicated cells were treated with ABT/S6/zVAD for the indicated intervals ± VbP. (**E**) Redox-state analysis of the indicated proteins in RAW264.7 cells following ABT/S6/zVAD treatment for the indicated times. Arrowheads indicate the oxidized and reduced protein species and the corresponding MIA40–TRX1 conjugates, as indicated. All cells, except in (**B**) and the PEG-labeled samples in (**E**), were lysed in buffer containing 40 mM NEM and analyzed by immunoblotting under reducing or nonreducing conditions. Samples in (**B**) were lysed without NEM and analyzed by immunoblotting only under reducing conditions. In (**E**), the indicated samples were labeled with 15 mM PEG11 or PEG24 and analyzed under reducing conditions. (**G**) DPP8/9 and PREP activity in lysates from RAW264.7, HT-1080, and A549 cells following ABT/S6/zVAD treatment for the indicated times, measured using GP-AMC and zGP-AMC, respectively. Compound concentrations: ABT-263 (10 μM), S63845 (1 μM), VbP (10 μM), zVAD (50 μM), and VX-765 (50 μM). Compounds in (**B**) were used at 10 μM except bortezomib (500 nM) and bafilomycin A1 (100 nM). Immunoblots are representative of three or more independent experiments. Asterisks denote non-specific bands.

**Figure S5.**
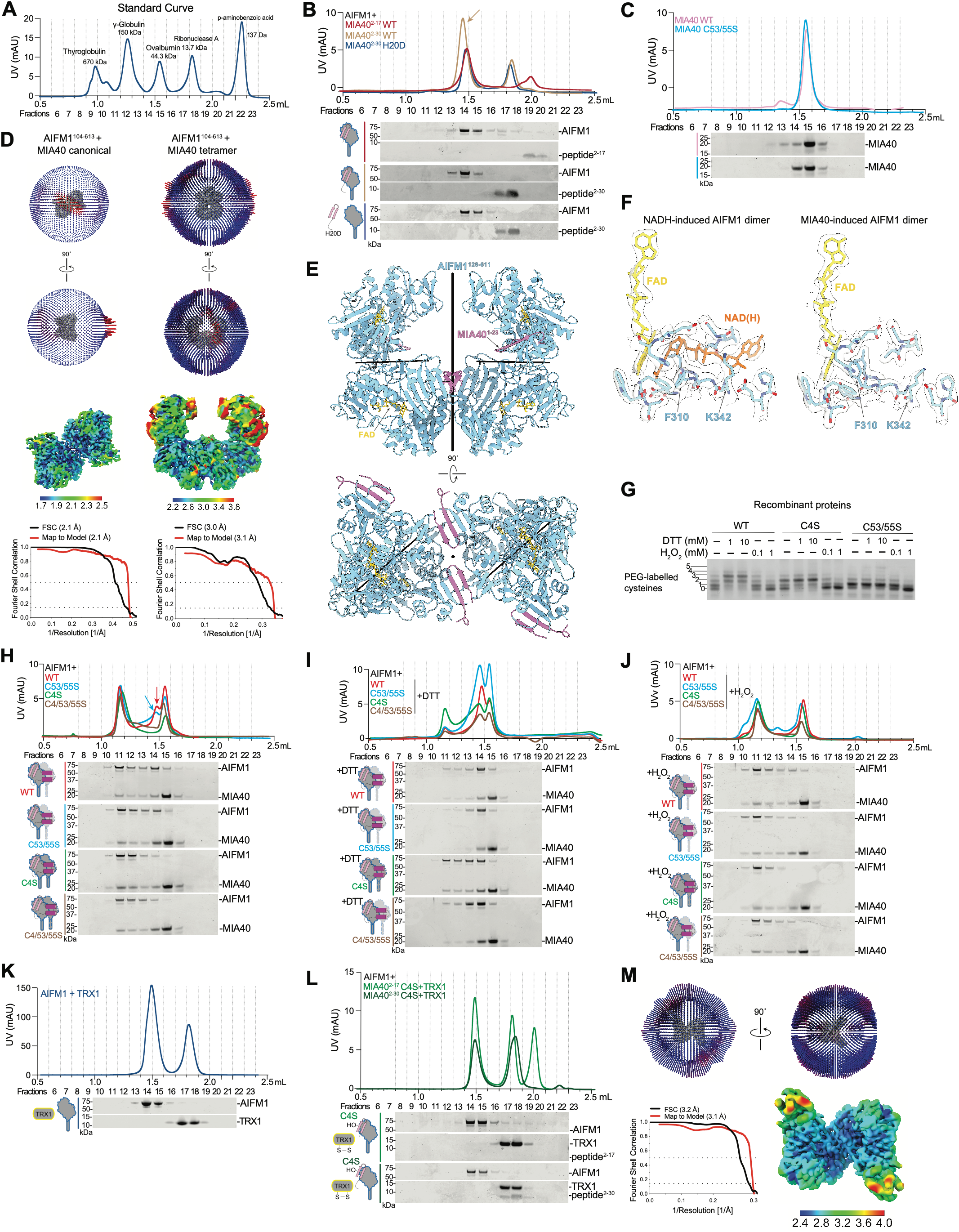
NADH-independent MIA40-driven AIFM1 dimerization. (**A**) SEC calibration using the indicated molecular weight standards. (**B**,**C**,**H-L**) SEC of the indicated proteins. DTT (1 mM) and H_2_O_2_ (100 μM) were added as indicated. The collected fractions were analyzed by Coomassie staining. Recombinant proteins were mixed at an AIFM1:MIA40:TRX1:peptide molar ratio of 1:2:4:10, with AIFM1 at 10 μM, except in (**K**) where AIFM1 is 35 uM. Arrow in (**B**) indicates binding of peptide to AIFM1 and arrow in (**H**) indicates the MIA40-bound AIFM1 monomer. TRX1 used in these experiments had Cys 62/69/73 mutated to Ser. In (**C**) the MIA40 WT chromatogram is also shown in Main **Fig. 5A**. (**D**,**M**) Cryo-EM particle orientation distributions, Fourier shell correlation curves, and cryo-EM maps of the canonical AIFM1–MIA40 dimer (left) and tetramer (right) reconstructions (**D**) and of the AIFM1–MIA40 peptide^2–17^–TRX1 complex (**M**). (**E**) Cryo-EM structure of the AIFM1–MIA40 tetramer (PDB 38EC). Thick and thin black lines indicate global and local twofold symmetry axes, respectively. (**F**) Comparison of cryo-EM density at the AIFM1 nucleotide-binding pocket between the structure in **Figure 1H** (left) and the current structure (right). (**G**) PEG24-maleimide analysis of purified MIA40 before and after DTT or H_2_O_2_ treatment at the indicated concentrations. Chromatograms are representative of three or more independent experiments.

**Figure S6.**
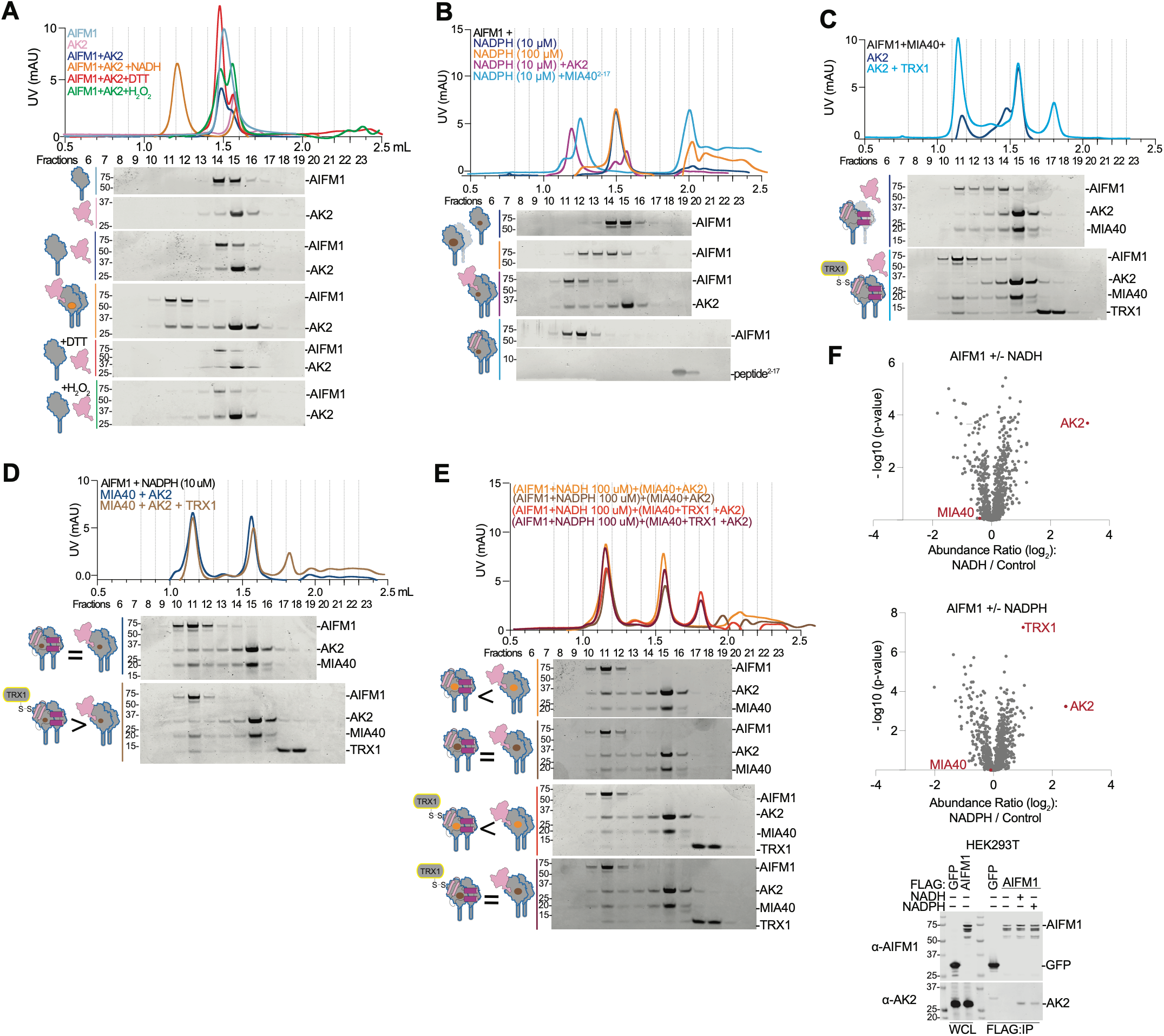
TRX1 increases stability of the MIA40-induced AIFM1 dimer. (**A-E**) SEC of recombinant AIFM1 with the indicated proteins, and peptides. DTT (1 mM), NADH (10 μM in **A** or as indicated), NADPH (10 or 100 μM as indicated) were added where indicated. Collected fractions were analyzed by Coomassie staining. Recombinant proteins were mixed at an AIFM1:MIA40:AK2:TRX1:MIA40 peptide molar ratio of 1:2:2:4:10, with AIFM1 at 10 μM. TRX1 Cys 62/69/73 Ser was used in these experiments. In (**A**), the AIFM1 alone chromatogram/blot is also shown in **Figure 5A**. (**F**) Quantitative mass spectrometry of FLAG–AIFM1 immunoprecipitates in the absence or presence of NADH or NADPH (1 mM), with immunoblot validation shown at right.

**Figure S7.**
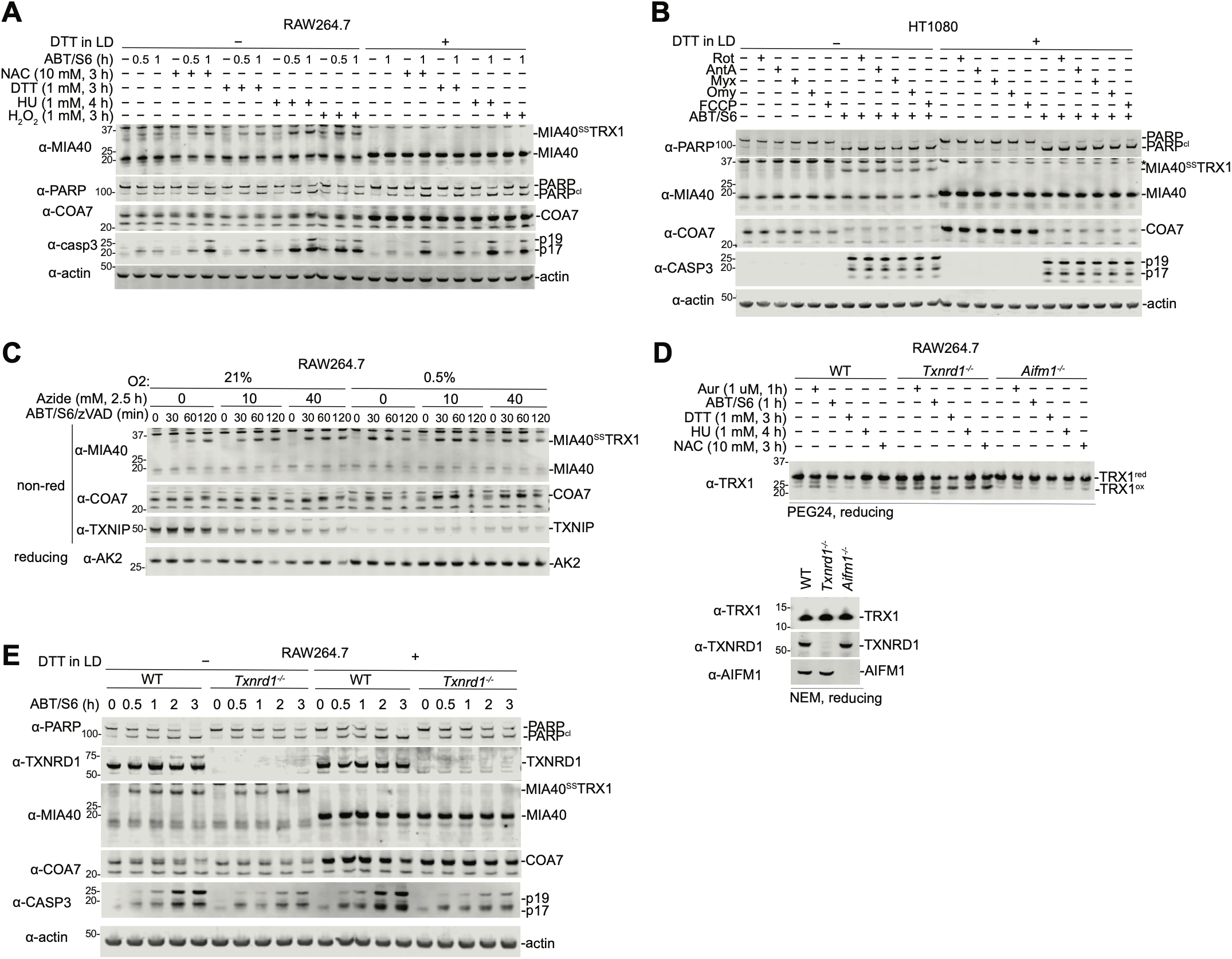
Redox perturbations do not alter conjugate formation. (**A–E**) The indicated cells were co-treated with ABT/S6 and the specified compounds for the indicated intervals. In (**C**), cells were put in hypoxia for 30 h prior to treatment with azide and ABT/S6 as indicated. All cells were lysed in 40 mM NEM, except panel (**D**, top), which was lysed in NEM PEG24 (15 mM). Samples were analyzed by immunoblotting under reducing or nonreducing conditions as indicated. Immunoblots are representative of three or more independent experiments.

